# Fluorescent Biosensors for Cryogenic Correlative Light and Electron Microscopy

**DOI:** 10.64898/2026.10.15.759416

**Authors:** Christopher A. Azaldegui, Kevin M. Rose, Davis Perez, Jacob A. Summers, Claudia A. Mak, William P. Dwyer, Anthony G. Vecchiarelli, Julie S. Biteen, José R. Dinneny, Peter D. Dahlberg

## Abstract

Cryogenic electron tomography provides nanometer-scale views of subcellular structures but lacks essential physiological context, such as the metabolic state or ion concentration. To bridge this gap, we incorporated fluorescent biosensors into a cryogenic correlative light and electron microscopy workflow, enabling direct measurement of physiological conditions alongside structural data. Here, we focused on establishing and calibrating biosensors for calcium, pH, and osmotic stress at cryogenic conditions through an *in vitro* workflow in which biosensors were frozen in defined chemical environments and then imaged ratiometrically. We then applied these biosensors across a range of sample types: an osmotic stress sensor in *Saccharomyces cerevisiae*, a calcium sensor in human U2OS cells, and a pH sensor in the bacterium *Halothiobacillus neapolitanus*. This approach links localized physiological changes to the 3D structures of macromolecules and their organization *in situ*.

## Introduction

Cryogenic electron tomography (cryoET) is a powerful tool for visualizing subcellular ultrastructure and large molecular complexes within their native cellular context (Förster and Briegel, 2024; Majumder and Zhang, 2025). However, cryoET alone does not provide information about the local physicochemical environment—such as metabolic state, pH, or ion concentrations—all of which influence the observed biological structures. While fluorescence microscopy is conventionally used in cryogenic correlative light and electron microscopy (cryoCLEM) workflows to target structures for milling or to localize proteins in tomograms (Dahlberg and Moerner, 2021; Pierson et al., 2024), fluorophores can also report on their environment and the physiological state of cells and organelles through changes in their photophysical properties (Greenwald et al., 2018; Frei et al., 2024). These fluorescent biosensors can be used to bridge the gap between structural biology and cellular physiology. We present a workflow to validate and apply genetically encoded fluorescent biosensors in cryoCLEM experiments.

Genetically encoded fluorescent biosensors enable the detection and measurement of a wide range of analytes and cellular events, including small molecules such as metabolites, ions, and secondary messengers (e.g., Ca^2+^, cAMP, ATP), as well as protein-protein interactions, post-translational modifications, and enzyme activities (Greenwald et al., 2018; Frei et al., 2024). Architecturally, most protein-based biosensors consist of one or more fluorescent proteins (FPs) fused to a sensing unit that undergoes a conformational change upon analyte binding, which is then transduced into an optical signal. Several readout mechanisms exist: single-FP intensiometric sensors report analyte binding as a change in fluorescence intensity driven by alterations to the local chemical environment of the chromophore. Other single-FP sensors also exploit dual excitation or emission peaks to achieve ratiometric readouts. Dual-FP sensors take two general forms: those that pair a responsive FP with a spectrally distinct and consistently bright FP as a normalization reference, correcting for variation in sensor concentration and illumination, and those based on Förster resonance energy transfer (FRET), in which analyte binding or changes in the physical or chemical environment brings two FPs into or out of proximity, producing an inherently ratiometric shift in donor-to-acceptor emission.

Despite their utility, only a handful of studies have reported the integration of fluorescent biosensors into cryoCLEM workflows. A few notable examples highlight the potential applications of biosensors in cryogenic samples. The diakylcarbocanine DiD dye was used to distinguish the fusion state of lipid bilayers *in vitro* (Metskas and Briggs, 2019) and the genetically encoded sensor, GCaMP, was used to identify calcium-dependent membrane contact sites in *Saccharomyces cerevisiae* (Hoffmann et al., 2019). We previously demonstrated that roGFP2 is a suitable ratiometric redox-state reporter at cryogenic temperatures and used it in a model bacterium to identify cell-cycle state (Perez et al., 2022).

Although many biosensors are available for room-temperature studies, not all are well suited for cryoCLEM. Photophysical state transitions, which require intramolecular rearrangements such as cis-trans isomerization, are often suppressed at cryogenic temperatures (Chang et al., 2014; Dahlberg et al., 2018). Therefore, we hypothesize that each sensor molecule is locked in the state it occupied at the moment of freezing, which can be read out after vitrification. Biosensors compatible with cryoCLEM must therefore meet several requirements. First, their molecular arrangement and corresponding spectral state must remain faithful to their environment after freezing. Second, because cryoCLEM captures a single snapshot of the sensor state at the moment of vitrification, the sensor value must be recoverable from that static measurement. This constraint makes intensiometric biosensors challenging to use, as their signal depends on both analyte level and local sensor concentration (Greenwald et al., 2018). Disentangling these two contributions normally requires time-dependent changes in signal due to the rapid changes in analyte concentration, which do not occur in frozen samples. Ratiometric sensors circumvent this problem because the ratio between two spectrally distinct states is intrinsically normalized against sensor concentration and remains informative even from a static measurement. However, because only a single time point is captured, the observable cannot be estimated by following an individual sensor molecule over time as it transitions between states. Instead, a robust physiological measurement requires an ensemble readout from many biosensors, typically within a diffraction-limited volume.

Here, we describe a generalizable experimental workflow to identify, validate, and calibrate biosensors for cryoCLEM. This workflow begins with the selection and *in vitro* validation of candidate sensors, followed by the detailed calibration of their spectral responses and, finally, implementation in biological systems for *in situ* studies. Using this approach, we validated and calibrated sensors for osmotic stress, Ca^2+^ concentration, and pH and demonstrated their utility for cryoCLEM applications across diverse biological systems. Through this process, we establish guiding principles for selection of fluorescent biosensors likely to function at cryogenic temperatures.

## Results

### Experimental workflow for the validation of fluorescent biosensors in cryoCLEM

Given the altered photophysical properties of fluorophores at cryogenic temperatures (Moerner and Orrit, 1999; Creemers et al., 2000; Regis Faro et al., 2011; Mantovanelli, 2023; Perez et al., 2024), fluorescent biosensors must be validated under cryogenic conditions to confirm that they retain their sensing properties (Dahlberg et al., 2018; Perez et al., 2022). In this experimental workflow, we first expose a purified biosensor to two defined chemical environments corresponding to the extremes of the analyte concentration range to which the biosensor responds. We then collect spectra from each sample at both ambient and cryogenic temperatures (Fig. 1a). This first experiment is intended to quickly validate the sensor at room temperature and exclude any sensor with minimal response under cryogenic conditions. If the response appears robust, we then calibrate the sensors by placing them in a series of defined chemical environments that vary only in analyte concentration before freezing the solutions. To mitigate artifacts from the air-water interface that may occur in thin protein samples produced by plunge-freezing on grids, we instead high-pressure freeze the calibration solutions to form a thick layer (100-200 µm) of vitrified protein at low micromolar concentrations (< 20 µM) (Fig. 1b). We then image these samples using the same cryogenic light microscope system intended for cellular imaging (Fig. 1c). The resulting ratiometric values provide a quantitative reference calibration curve for cryoCLEM of cells, mapping spectral ratio to analyte concentration.

**Figure 1:**
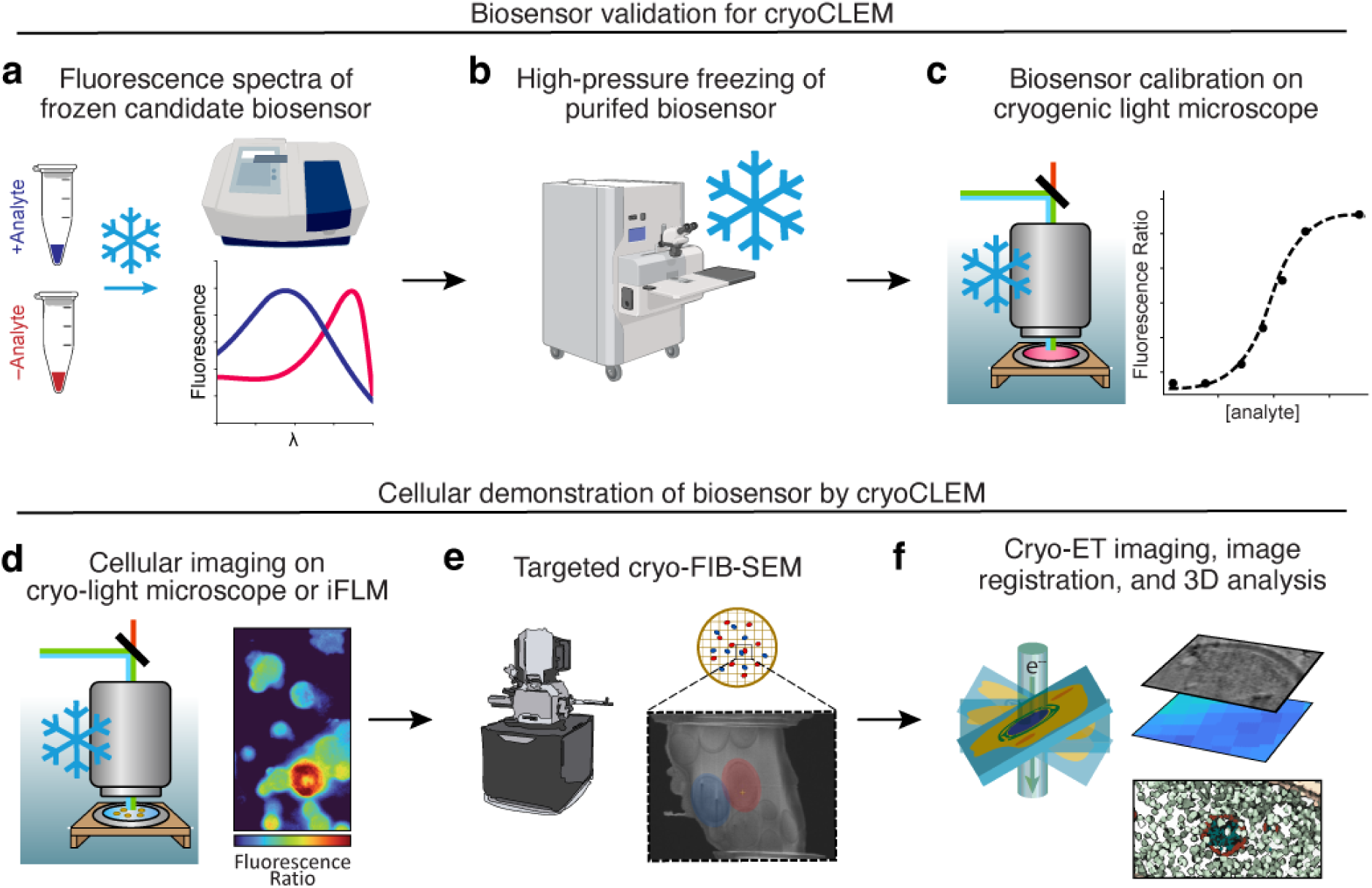
Biosensor cryoCLEM Workflow. **a)** Biosensors are quickly screened for viability by collecting fluorescence spectra in the presence of different analyte concentrations under cryogenic conditions. **b**) Candidate biosensors are frozen in calibration standards in a high-pressure freezer. **c**) HPF-prepared calibration standards are ratiometrically imaged on a cryogenic light microscope. **d**) Ratiometric images of frozen cellular samples are collected on a cryogenic light microscope. **e**) Ratiometric images are used for targeting during cryo-FIB milling. **f**) Tilt series are collected and subsequently reconstructed and registered with the ratiometric data.

Following validation and calibration, we perform cryoCLEM on cellular samples. We conduct ratiometric imaging of the biosensor expressed in vitrified cells using the same optical microscope and imaging conditions as the calibrations to ensure quantitative measurement of the target analyte (Fig. 1d). The resulting ratiometric image guides the selection of specific cells for lamella preparation by focused ion beam (FIB) milling; when calibration is performed using microscopes integrated into the FIB-milling system, the final lamella can also be imaged ratiometrically (Fig. 1e). Lastly, we perform tomography, 3D analysis, and image registration, connecting the structural data with the physiological context (Fig. 1f).

### SED1 is an osmotic stress sensor suitable for cryoCLEM

First, we sought to identify an osmotic stress sensor compatible with cryoCLEM because osmotic fluctuations are a common physicochemical change that cells experience. These fluctuations stem from intrinsic factors, such as cell-cycle-dependent metabolite shifts or ion channel activity (Yancey, 2005; Khatibi et al., 2019). For instance, decreased external water content drives efflux, increasing intracellular solute concentration, molecular crowding, and viscosity (Record et al., 1998; Spitzer and Poolman, 2009; Sukenik et al., 2017). Sensor Expressing Disorder Protein 1 (SED1) is a FRET-based sensor comprising a cyan fluorescent protein (CFP) donor (mCerulean3), a yellow fluorescent protein (YFP) acceptor (Citrine), and an osmotically sensitive intrinsically disordered linker (AtLEA4-5) (Figure 2a) (Cuevas-Velazquez et al., 2021). In high-salt or crowded environments, the AtLEA4-5 linker folds, bringing the fluorophores into proximity, thereby increasing FRET efficiency and producing a ratiometric response.

**Figure 2:**
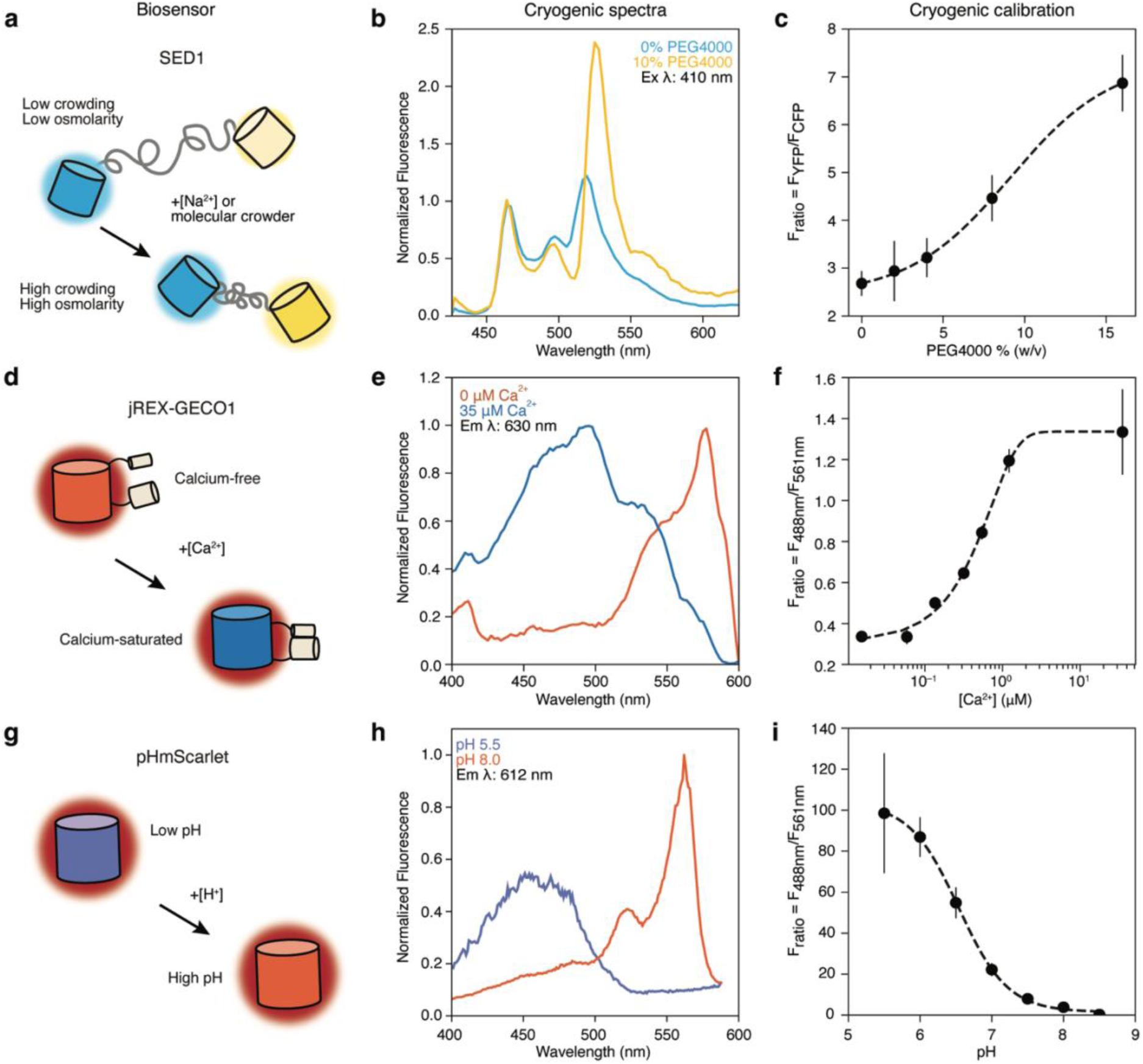
Biosensors for cryoCLEM. **a)** SED1 osmotic stress mechanism. FRET pairs mCerulean (cyan) and Citrine (yellow) are linked by an intrinsically disordered protein that is sensitive to osmotic stress. **b)** Cryogenic fluorescence emission spectra of SED1 in the presence and absence of 10% PEG4000 (w/v). Emission spectra are normalized to their respective mCerulean peak at 465 nm. **c)** Fluorescence ratio (F_ratio_) of Citrine (YFP) and mCerulean (CFP) emission of high-pressure frozen SED1 as a function of PEG4000 concentration. **d)** jREX-GECO1 calcium sensing mechanism. A calmodulin-based (gray) conformational change shifts the chromophore’s absorption maximum from 570 to 485 nm in the presence of calcium. **e)** Cryogenic fluorescence excitation spectra of jREX-GECO1 in the presence and absence of Ca^2+^ (35 µM). Excitation spectra are normalized independently. **f)** The F_ratio_ of 488- and 561-nm excitation of HPF jREX-GECO1 as a function of Ca^2+^ concentration. **g)** Deprotonation of the pHmScarlet chromophore red shifts and increases its absorbance. **h)** Cryogenic fluorescence excitation spectra of pHmScarlet at pH 5.5 and pH 8. Excitation spectra are normalized to the maximum of the pH 8 curve. **i)** The F_ratio_ of 488- and 561-nm excitation of high-pressure frozen pHmScarlet as a function of pH. All calibration curve points are the average of at least five ratiometric measurements for each condition. The error bars represent the standard deviation. The dashed lines are the fit to a continuous logistic function.

We validated the response of purified SED1 at room temperature (RT) and 77 K using 0% and 10% PEG4000 as a synthetic crowder. In the absence of the crowder, the YFP emission peak at 520 nm is approximately 80% of the CFP emission peak at 470 nm (Supplementary Fig. 1a). Addition of the crowder increased the YFP emission peak to roughly 100% of the CFP emission peak (Supplementary Fig. 1a), consistent with an increase in FRET. At 77 K, in the absence and presence of crowder, the YFP emission peak increased relative to RT (Supplementary Fig. 1b-c) indicating an increased FRET efficiency at 77 K. Despite this effect, the YFP emission peak increased from 120% to over 200% of its corresponding CFP emission peak as a result of the crowder (Fig. 2b). This result demonstrates that SED1 remains sensitive to molecular crowding at 77 K and that this sensitivity is enhanced over 4-fold compared to room temperature.

To calibrate SED1 at 77 K, we high-pressure froze the purified sensor in solutions with various PEG4000 concentrations and loaded them onto a cryogenic light microscope (Supplementary Fig. 1d-e). Consistent with previous results (Cuevas-Velazquez et al., 2021), the F_ratio_ of SED1 increased in a PEG4000 concentration-dependent manner and exhibited a 3-fold increase in F_ratio_ (YFP/CFP Em) from 0 to 12% PEG4000 (Fig. 2c).

### jREX-GECO1 is a calcium biosensor suitable for cryoCLEM

Given the critical role of calcium in cell signaling (Berridge et al., 2000; Giorgi et al., 2018a; Giorgi et al., 2018b), we sought to identify a calcium sensor suitable for cryogenic studies. We first tested MatryoshCaMP6s, a GCaMP-based sensor that uses a large Stokes shift (LSS) mOrange as a covalently linked reference reporter (Supplementary Fig. 2) (Ast et al., 2017). The two fluorophores exhibit overlapping excitation spectra at 440 nm and emit at 510 nm (GCaMP6s) and 570 nm (LSSmOrange), enabling an emission-based ratiometric readout. At RT and in the absence of Ca^2+^, the emission of GCaMP6s at 510 nm peaks at 15% of the 570 nm reference peak of LSSmOrange (Supplementary Fig. 2a). Upon Ca^2+^ binding, GCaMP6s emission increased significantly to 115% of the reference peak (Supplementary Fig. 2a). Intriguingly, at 77 K, GCaMP6s is largely emissive even in the absence of Ca^2+^, with its 510 nm peak reaching 85% of the reference peak at 570 nm. This result indicates that at 77 K, the presence of a high Ca^2+^ concentration is not necessary to relieve the fluorescence quenching of the GCaMP6s chromophore. Indeed, a previous report on the GFP chromophore described that steric barriers to the chromophore rotation enable fluorescence in otherwise non-fluorescent, unconstrained environments (Maddalo and Zimmer, 2006). At 77 K, the chromophore in GCaMP6s is presumably rigidified, even in its open state, resulting in fluorescence that makes it unsuitable for further cryoCLEM experiments. This negative result highlights the importance of the validation and calibration of biosensors under cryogenic conditions prior to their use in correlative experiments.

We next tested the red calcium biosensor jREX-GECO1 (Fig. 2d), a variant of the large Stokes shift REX-GECO1 sensor (Molina et al., 2019; Wu et al., 2014). Between its Ca^2+^-free and Ca^2+^-bound states, jREX-GECO1 shifts its excitation maximum from ∼575 nm to ∼480 nm while maintaining red emission (∼585–600 nm) (Supplementary Fig. 3a). These spectral shifts are maintained at cryogenic temperatures (Fig. 2e and Supplementary Fig. 3b-c). Calibration of high-pressure-frozen jREX-GECO1 in a wide range of Ca^2+^ concentrations highlighted its dynamic range of 100 nM to approximately 2 µM, resulting in a nearly 4-fold increase in F_ratio_ across this range (Fig. 2f and Supplemental Fig. 3d-e). The typical average resting Ca^2+^ concentration in cells is approximately 100 nM (Berridge et al., 2000), making jREX-GECO1 an ideal sensor to detect intracellular calcium spikes in cryoCLEM experiments.

### pHmScarlet is a robust ratiometric biosensor at cryogenic temperatures

Intracellular pH modulates protein structure and function, regulating membrane transport, energy production, and organelle homeostasis (Casey et al., 2010). Although typically buffered to a pH of 7.2–7.4, cells exhibit pH fluctuations during the cell cycle or in response to external stimuli. In eukaryotes, altered pH levels often correlate with specific disease states (Obara et al., 2008; Vaughan-Jones et al., 2009; Pérez-Sayáns et al., 2009). Eukaryotes and prokaryotes also compartmentalize different chemical environments within organelles. For example, lysosomes in eukaryotes contain a pH < 5; similarly, carbon-fixing bacterial microcompartments are proposed to have a more acidic internal environment compared to the surrounding cytoplasm (Menon et al., 2010; Mangan et al., 2014; Huang et al., 2022). Given the central roles of pH in cellular and organelle homeostasis, we explored the applicability of pH sensors for cryoCLEM.

We evaluated pHRed and pHluorin2, two excitation-based ratiometric pH sensors with red and green emission, respectively (Tantama et al., 2011; Mahon, 2011), as well as pHmScarlet, an intensity-based pH sensor (Liu et al., 2021). At RT and pH 7.5, pHluorin2 excitation peaks at 390 nm, whereas at pH 5.5, it shifts to 470 nm (Supplemental Fig. 4a). At 77 K, the pH-7.5 excitation peak shifts to 400 nm, while at pH 5.5 its peak shifts to 460 nm (Supplemental Fig. 4b). Similarly, the excitation spectrum of pHRed at pH 5.5 peaks at 580 nm, while at pH 9.0 another peak emerges at 430 nm (Supplemental Fig. 4c). At 77 K, both excitation peaks narrow and the 430-nm peak shifts to 415 nm (Supplemental Fig. 4d). Despite these slight shifts in peak wavelength, the ratiometric response of pHluorin2 and pHRed at 77 K to changes in pH is detectable and comparable to room-temperature.

While primarily used as an intensiometric sensor, pHmScarlet (Fig. 2g) exhibits an absorbance shift from 560 nm to 500 nm below pH 7, suggesting potential for ratiometric readout (Liu et al., 2021). Consistent with previous reports, RT pHmScarlet showed poor excitation at pH 5.5—with a small peak at 460 nm—and a prominent excitation peak at 560 nm at pH 8 (Supplementary Fig. 5a). At 77 K and pH 8, excitation maxima remained consistent; however, the 460 nm excitation peak at pH 5.5 increased more than ten-fold relative to RT (Figure 2h and Supplemental Fig. 5b-c). We hypothesize that the loss of non-radiative decay pathways at 77 K enables the pHmScarlet chromophore to undergo efficient excited-state intramolecular proton transfer (ESIPT) (Zhou and Han, 2018), resulting in its red emission. This enhancement in emission at cryogenic temperatures made pHmScarlet an effective ratiometric sensor at 77 K; we therefore selected it and pHRed for further calibration. We excluded pHluorin2 to prioritize red-emitting sensors, thereby avoiding the high background autofluorescence typical of green wavelengths in cryogenically prepared cellular samples (Carter et al., 2018; Perez et al., 2022); however, it is likely that pHluorin2 could also function in cryoCLEM workflows once calibrated if a green sensor were desired.

We evaluated the response of high-pressure frozen pHRed and pHmScarlet across a range of pH values to determine the optimal pH sensor for cryoCLEM. However, imaging of pHRed under 561 nm excitation resulted in rapid photobleaching (Supplementary Fig. 6a), indicating poor photostability at 77 K. Although reducing laser power enabled an F_ratio_ measurement, which yielded a 4.6-fold increase across pH 5.5 to 9 (Supplemental Fig. 6b), the poor photostability limits the utility of pHRed for cryoCLEM where weaker *in situ* fluorescence measurements are likely to be perturbed by the measurement itself. This photosensitivity is another potential failure mode of biosensors under cryogenic conditions and should be evaluated during calibration. In contrast, pHmScarlet exhibited no such issues and a wide dynamic range (Supplemental Fig. 6c-d and Fig. 2i), with a 300-fold reduction in F_ratio_ from pH 5.5 to 8.5. These findings establish pHmScarlet as a superior ratiometric sensor for measuring intracellular pH at cryogenic temperatures.

Following the validation and calibration of these biosensors (SED1, jREX-GECO1, and pHmScarlet), we next demonstrated their utility in cellular cryoCLEM experiments. Here, we aimed to show the versatility of the approach by performing experiments with the identified biosensors across three biological model systems and using different optical systems.

### SED1 correlates osmotic stress and its ultrastructural effects in S. cerevisiae

Hyperosmotic shock draws water out of the cell, reducing membrane tension and causing cell shrinkage (Sukenik et al., 2017; Colom et al., 2018). This reduction of membrane tension results in membrane invaginations and vesicle formation (Phan et al., 2025). As a first proof-of-principle experiment, we expressed SED1 (molecular crowding sensor) cytoplasmically in *S. cerevisiae* to correlate hyperosmotic stress with these membrane deformations. We treated cells with 1 M NaCl for 10 min or left them untreated before plunge-freezing (Fig. 3a). We then performed cryogenic light microscopy on these samples using the same optical setup as the SED1 calibration. Cryogenic ratiometric analysis showed that while untreated cells exhibited a low F_ratio_ (1.63 ± 0.25), the osmotically stressed cells displayed a higher average F_ratio_ (3.60 ± 1.54) and a significantly larger standard deviation (Fig. 3b-c and Supplementary Fig. 7a-b), demonstrating the elevated molecular crowding and highlighting the cell-to-cell heterogeneity in osmotic stress.

**Figure 3:**
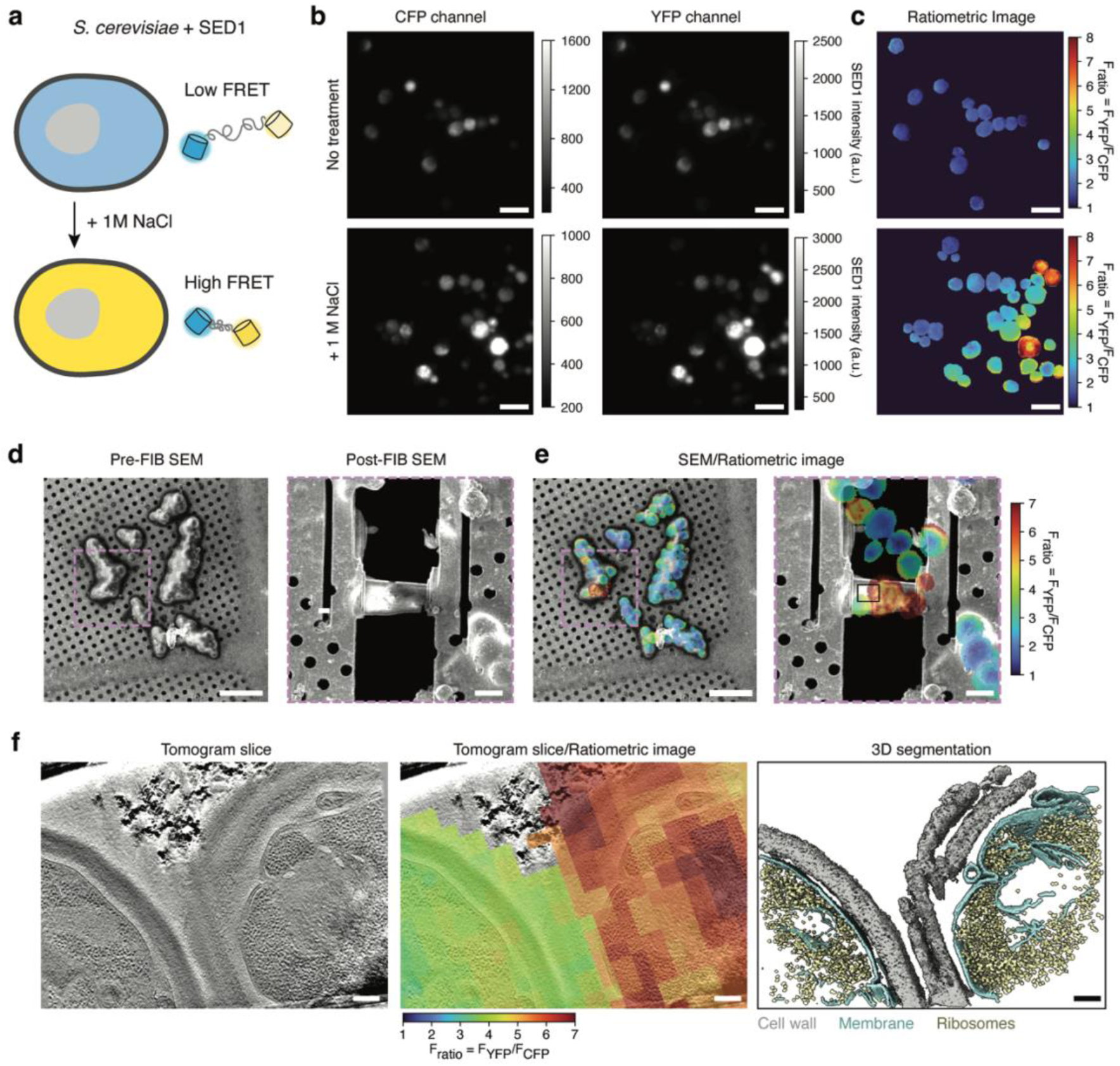
SED1 cryoCLEM of Osmotically Stressed *S. cerevisiae* Cells. **a)** Diagram of osmotic stress experiment in *S. cerevisiae*. **b)** Fluorescence images of untreated and osmotically stressed (+1 M NaCl) cells collected using 465/30 nm (CFP) and 545/40 nm (YFP) bandpass filters. **c)** Corresponding F_ratio_ images from **b**. Scale bars: 10 µm. **d)** SEM image of NaCl-treated cells. Scale bar: 20 µm. Inset: Post-FIB-milling image of lamella site. Scale bar: 5 µm. **e)** As in **d** but with a registered ratiometric image depicting two adjacent cells with different F_ratio_ signal. Black inset indicates the tomogram region shown in **f**. **f)** Left shows a slice from the reconstructed tomogram collected from the lamella in **d-e**. The middle shows the tomogram slice overlaid with the registered F_ratio_ signal. The right shows the segmentation of ribosomes (gold), membrane (teal), and cell wall (gray). Scale bars: 200 nm.

To determine whether variations in osmotic stress correlate with changes in ultrastructure, we prepared treated *S. cerevisiae* cells for cryoET. We targeted cells with a broad range of F_ratio_ values for focused ion beam (FIB) milling (Fig. 3d-e). Registering the ratiometric image onto the reconstructed tomogram allowed us to visualize the ultrastructural differences that were directly correlated with osmotic stress (Fig. 3f and Supplemental Fig. 7c-d). For example, cells with moderate osmotic stress levels (F_ratio_ < 5, Fig. 3f yellow cell) maintained an intact cell wall and exhibited only minor plasma membrane invaginations (Fig. 3f and Supplemental Fig. 7d). In contrast, cells with a high osmotic-stress signal (F_ratio_ > 5, Fig. 3f orange cell) displayed striking cell wall deformations, extensive detachment of the plasma membrane, and vesicle formation. This deformation of the plasma membrane is consistent with cellular volume decreases observed in cells exposed to hypertonic solutions (Roffay et al., 2021; Liu et al., 2023; Phan et al., 2025).

### CryoCLEM of calcium influx in U2OS cells with jREX-GECO1

To demonstrate jREX-GECO1 (Ca^2+^ sensor) in a cryoCLEM experiment, we expressed it in U2OS cells to monitor the ultrastructural effects of a rapid influx of extracellular Ca^2+^. Rapid calcium release from the endoplasmic reticulum into the cytosol can lead to the activation of mitochondrial-mediated apoptotic pathways, resulting in mitochondrial depolarization and matrix swelling (Giorgi et al., 2018a). To artificially induce this effect, we treated cells with the pore-forming toxin Streptolysin O (SLO) in a high-Ca^2+^ medium, producing a Ca^2+^ influx prior to plunge-freezing (Fig. 4a). To highlight the accessibility of this approach, we performed cryogenic sensor calibration and ratiometric imaging using the fluorescence light microscope (iFLM) system integrated into the Aquilos2 cryo-FIB-SEM prior to lamella preparation. Because jREX-GECO1 is an excitation-ratiometric sensor, we collected emission during sequential illumination at 470 nm and 565 nm (Fig. 4b). While untreated cells displayed higher signals under 565 nm excitation—consistent with jREX-GECO1 being in its Ca^2+^-free state—SLO-treated cells showed a marked signal increase under 470 nm excitation.

**Figure 4:**
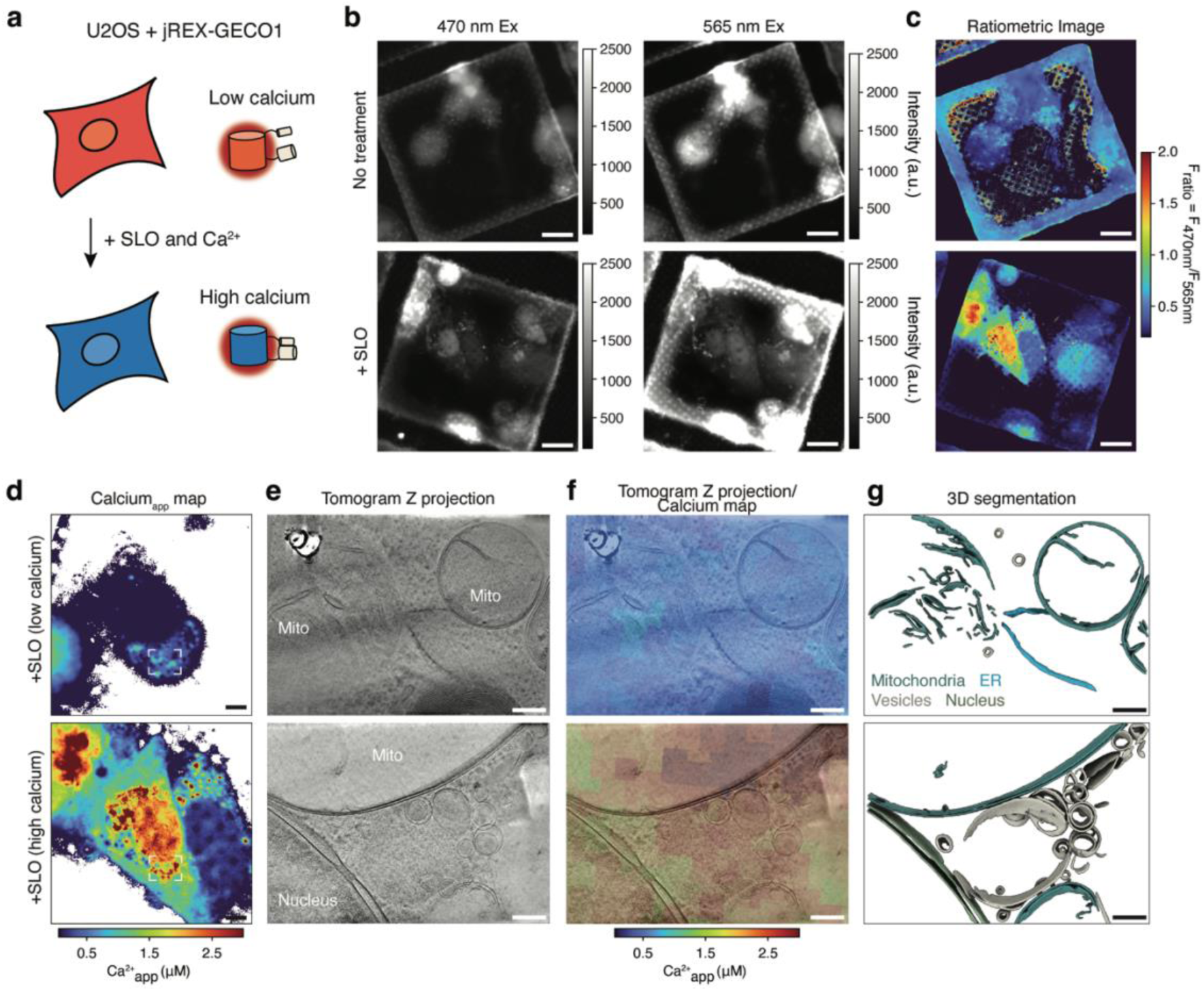
jREX-GECO1 cryoCLEM of Calcium Influx in U2OS Cells. **a)** Schematic of calcium spike experiment in U2OS cells. **b)** Fluorescence images of untreated and SLO-treated cells collected using 470 nm and 565 nm excitation sources. **c)** Ratiometric image, F_470nm_/F_565 nm_, of fluorescence channels in **b**. Scale bars: 20 µm. **d)** Calcium maps of SLO-treated U2OS cells with low and high Ca^2+^ concentrations. White dashed box shows the region of the cell where a lamella was generated. Scale bars: 5 µm. **e)** Reconstructed tomogram z projection of the region within the cells shown in **d**. **f)** Reconstructed tomogram z projection overlaid with the correlated calcium map. **g)** Segmentation of mitochondria (teal), ER (blue), vesicles (gray), and the nuclear membrane (green). Scale bars: 200 nm.

We applied a calibration model also collected on the iFLM (Supplementary Fig. 8a) to these ratiometric data (Fig. 4c) to generate an apparent Ca^2+^ concentration (Ca^2+^) map, which revealed elevated Ca^2+^ concentration in SLO-treated cells (Fig. 4d and Supplementary Fig. 8b). We then collected tomograms of untreated and SLO-treated cells and observed differences in mitochondrial ultrastructure. In untreated cells, mitochondria were fragmented but maintained intact cristae (Supplementary Fig. 8c), whereas SLO-treated cells contained enlarged, swollen mitochondria characterized by the loss of cristae (Fig. 4e-g and Supplementary Fig. 8d-e). Correlation with the Ca^2+^_app_ maps revealed that this ultrastructural defect is more pronounced in cells with higher Ca^2+^_app_ (Fig. 4f and Supplementary Fig. 8d-e). Indeed, in cells with a lower Ca^2+^_app_ (< 0.5 µM), mitochondria contained 4.7 ± 3.0 cristae, while mitochondria from cells with Ca^2+^_app_ > 0.5 µM contained 1.4 ± 0.7 cristae. Both values are strikingly lower than the cristae count in untreated cells (12.1 ± 2.7) and in previous reports (Rose et al., 2025). These phenotypes—mitochondrial swelling and cristae loss—align with features reported in apoptotic cells (Ader et al., 2019) and are consistent with mitochondrial failure triggered by elevated cytoplasmic Ca^2+^ (Hajnóczky et al., 2003).

### CryoCLEM of a carbon-fixing bacterial microcompartment using pHmScarlet

To demonstrate pHmScarlet in a cellular cryoCLEM experiment, we labeled carboxysomes—bacterial microcompartments that encapsulate RuBisCO and carbonic anhydrase to concentrate CO_2_ and optimize carbon fixation (Kerfeld et al., 2018). Because the activities of RuBisCO and carbonic anhydrase are pH-dependent, carboxysomes are thought to maintain a pH gradient relative to the cytoplasm.

Specifically, a proton gradient with a lower internal pH is proposed to drive the equilibrium of HCO_3_^−^ to CO_2_, facilitating its conversion to organic carbon by RuBisCO (Menon et al., 2010; Mangan et al., 2014). Given that carboxysomes are structurally heterogeneous, displaying variations in size, composition, and enzyme packing (Metskas et al., 2022; Ni et al., 2022), we utilized pHmScarlet to explore potential correlations between carboxysome structure and luminal pH. We fused pHmScarlet to the small subunit of RuBisCO (CbbS) to specifically target the sensor to the carboxysome lumen in *Halothiobacillus neapolitanus* cells (Fig. 5a).

**Figure 5:**
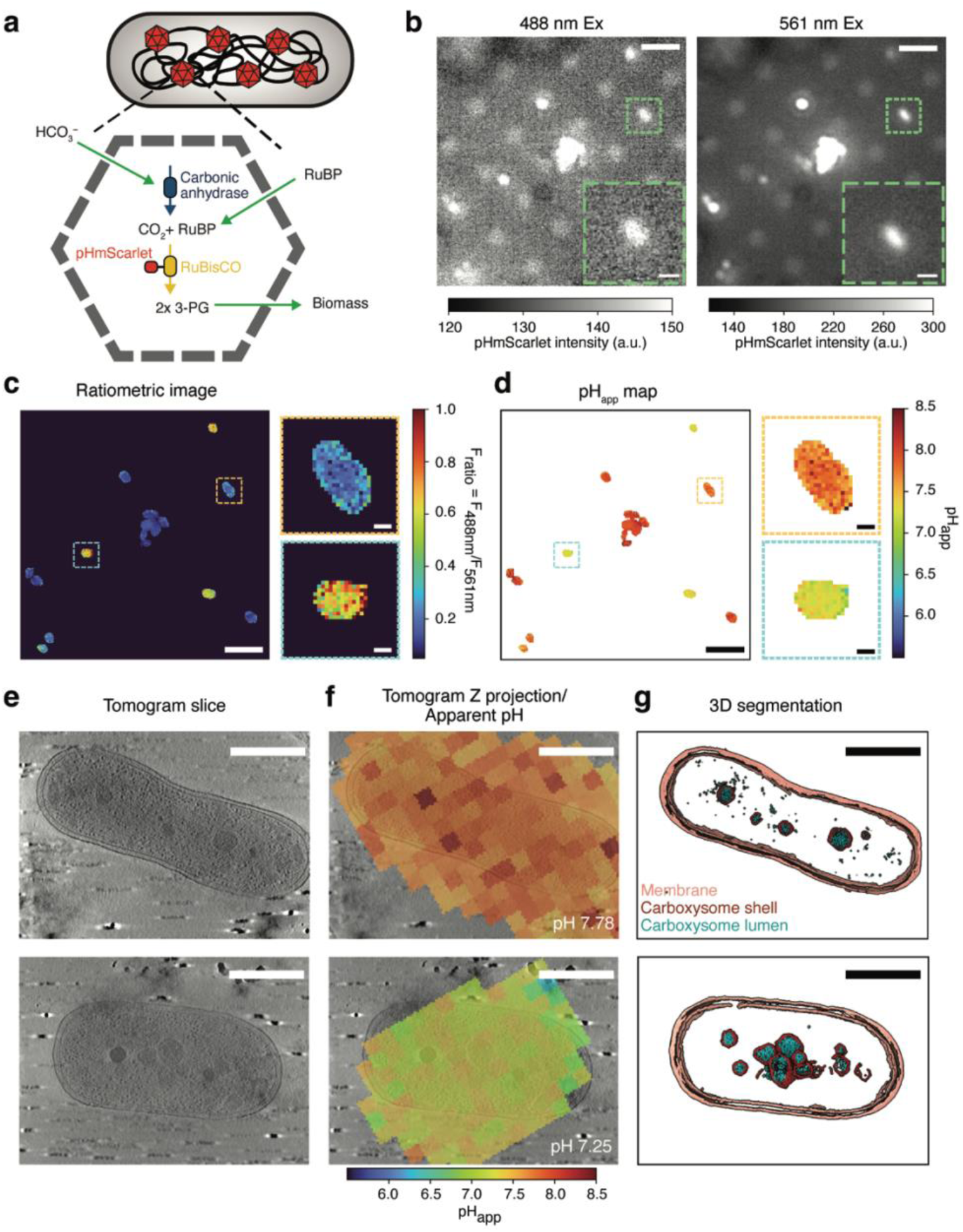
pHmScarlet cryoCLEM of Carboxysomes in *Halothiobacillus neapolitanus*. **a)** Schematic of carboxysome function, see text for details. **b)** Fluorescence images of *H. neapolitanus* cells collected using 488 nm and 561 nm excitation sources. Inset shows the fluorescent signal from a single cell. Scale bars: 5 and 1 µm. **c)** Ratiometric image, F_488 nm_/F_561 nm_, of fluorescence images shown in **b** after correcting for the background. Insets: Single cells with different F_ratio_. **d)** Apparent pH of cells in **c**. Insets: Single cells with different apparent pH. Scale bars: 5 µm and 500 nm. **e)** Reconstructed tomogram slice of the two cells highlighted in **c-d**. **f)** Registered pH image overlaid with tomogram slice from **e**. **g)** Segmentation of membrane (tan), ribosomes (green), carboxysome shell (red), and carboxysome lumen contents (teal). Scale bars: 500 nm.

We performed ratiometric imaging of plunge-frozen cells expressing CbbS-pHmScarlet (Fig. 5b-c). While our cryogenic light microscope cannot resolve individual carboxysomes because they are separated by less than the optical diffraction limit, the measurements provide a robust determination of their average ratiometric signal across single cells. Using our calibration model (Fig. 2i and Supplementary Fig. 9a), we converted the F_ratio_ images into apparent pH (pH_app_) maps (Fig. 5d). The resulting average single-cell pH distribution of 7.67 ± 0.26 (Supplementary Fig. 9b) is consistent with the pH measured in heterologously expressed α-carboxysomes in *Escherichia coli* (Huang et al., 2022), demonstrating the ability of pHmScarlet to accurately report intracellular pH at cryogenic temperatures. Correlation with whole-cell tomograms of *H. neapolitanus* shows distributed carboxysomes with an average pH of 7.78, whereas clustered carboxysomes show a lower average pH of 7.25 (Fig. 5e-g). While the data shown here are not sufficient to make claims about correlations between the chemical environment and structure or spatial distribution of carboxysomes, it highlights the potential for pHmScarlet and biosensor-based CryoCLEM as an approach to link carboxysome structure and physiology in future studies.

## Discussion

In this work, we established an experimental approach to validate and calibrate ratiometric fluorescent biosensors in correlative light and electron microscopy experiments. Using this workflow, we identified fluorescent biosensors compatible with cryogenic temperatures that report on molecular crowding, Ca^2+^ concentration, and pH. Furthermore, we demonstrated their use across different model biological systems: bacteria (*H. neapolitanus*), yeast (*S. cerevisiae*), and human cell lines (U2OS), highlighting the broad applicability of this method to directly correlate physiological states with cellular ultrastructure.

Temperature-dependent changes in biosensor photophysics must be accounted for in cryoCLEM workflows, motivating the need to characterize each sensor under cryogenic conditions. Our fully cryogenic validation and calibration procedure corrects for these effects, including the narrowing of excitation and emission spectra caused by suppression of thermal broadening at low temperature (Creemers et al., 2000). The procedure also accounts for changes in non-radiative decay pathways, many of which are suppressed at 77 K and can alter fluorophore quantum yields (Moerner and Orrit, 1999; Hulleman et al., 2018). Together, these photophysical effects may explain both the changes in FRET efficiency observed for SED1 and the efficient red emission of pHmScarlet under blue (460 nm) excitation (Fig. 2h), which is essential for its function as a ratiometric sensor at 77 K.

Our investigation establishes key principles for the selection and design of biosensors compatible with cryoCLEM. We demonstrate that FRET-based sensors, such as SED1, serve as a successful template for cryogenic studies. Their sensing mechanism relies on large-scale conformational changes, which are effectively trapped by vitrification, thereby preserving fluorescence signatures of the sensor in a particular physiological state. However, the isotropic averaging of dipole orientations that normally accompanies FRET measurements at room temperature is no longer valid at cryogenic temperatures. Instead, FRET readings must rely on the average of many sensors within a diffraction-limited volume for an accurate readout. Future work could push these measurements to the single-molecule regime if both the donor and acceptor dipole orientations could be determined at the single-molecule level. We also identified a biosensor, MatryoshCaMP6s, which fails to preserve its calcium-dependent response, highlighting a critical constraint for the implementation of biosensors for cryoCLEM. Sensors that modulate fluorescence through the dynamic flexibility of the chromophore environment lose their sensitivity at 77 K. In this scenario, the flexibility that is required to quench fluorescence is arrested by vitrification, locking the sensor in a bright state independent of its target analyte. Future work focused on the characterization of biosensor structure and photophysics under cryogenic conditions would aid in the rational design of biosensors for these applications.

Despite their value, fluorescent biosensors in cryoCLEM have limitations. First, our use of ratiometric biosensors requires two spectral channels, constraining multicolor imaging with other fluorescent markers. Second, because our cryogenic ratiometric imaging relies on the measurement of an ensemble of molecules in space rather than repeated measurements over time for an accurate environmental reading, the approach has limited spatial resolution. Consequently, our measurements in *S. cerevisiae* and U2OS cells were averaged at the single-cell level, and we could not resolve the pH of individual carboxysomes. Future advancements, such as the implementation of cryogenic structured illumination microscopy (cryo-SIM) (Phillips et al., 2020), could help overcome this resolution limit for sufficiently concentrated biosensors.

Although substantial work remains to further improve and generalize the method, the ability to directly correlate physiology and structure using biosensor-based cryoCLEM opens new avenues for cell and structural biology. By linking physiological state to ultrastructure, this approach can make structure–function relationships easier to establish, particularly in biological systems with substantial heterogeneity where the low-throughput nature of cryoET makes structural differences challenging to interpret. Biosensor-based cryoCLEM can also be combined with subtomogram averaging (Schur, 2019), enabling specific structures to be quantified *in situ* as a function of defined physiological variables. More broadly, this approach highlights the power of fluorescence microscopy to move beyond identifying the locations of specific biomolecules in cryoET workflows, instead providing a rich physiological context for interpreting the structures observed.

## Materials and methods

### Protein Purification

#### Purification of pHRed

Overexpression and purification of pHRed were adapted from Tantama et al (2011), with modifications. An overnight culture of DH5*αE. coli* cells expressing N-terminal His-tagged pHRed from a pRsetB-pHRed plasmid (AddGene, Cat#31472) was inoculated (1:100) into a 1 L culture of LB + carbenicillin (100 µg/mL) and grown at 37 °C with shaking until an OD600 of 1–1.2. Cells were pelleted and stored at -80 °C. Bacterial pellets were resuspended in 40 mL lysis buffer [100 mM MOPS pH 7.4; 50 mM KCl; 5 mM NaCl; 0.5 mM MgCl_2_; 20 mM Imidazole pH 7.4; 10% glycerol; 0.05 mg/mL Lysozyme (Thermo Fisher); protease inhibitor tablet (Thermo Fisher)], sonicated using cycles of 10 s on and 20 s off at 50% power for 6 min. Lysates were clarified via centrifugation at 13,000 rpm at 4 °C for 30 min. Clarified lysates were passed through a 0.45-µm filter and loaded onto a 1-mL HisTrap HP (Cytiva) equilibrated in buffer A [100 mM MOPS pH 7.4; 50 mM KCl; 5 mM NaCl; 0.5 mM MgCl_2_; 20 mM Imidazole pH 7.4; 10% glycerol; 5 mM 2-Mercaptoethanol (BME)]. Columns were washed with 5 column volumes of 5% buffer A. Elution was performed using a 5–100% gradient of buffer B [100 mM MOPS pH 7.4; 50 mM KCl; 5 mM NaCl; 0.5 mM MgCl_2_; 500 mM Imidazole pH 7.4; 10% glycerol; 5 mM BME] via an AKTA Pure system (Cytiva). Fractions with the highest yield were pooled, concentrated, passed through a 0.45-µm filter, followed by a size-exclusion column. Peak fractions were pooled, concentrated in storage buffer [10 mM HEPES pH 7.4; 120 mM KCl; 10 mM NaCl; 0.5 mM MgCl_2_, 10% glycerol], and stored at -80 °C.

#### Purification of SED1, MatryoshCaMP6s, jREX-GECO1, pHluorin2, and pHmScarlet

Purification of other fluorescent biosensors was performed according to Cranfill et al., with minor adjustments (Cranfill et al., 2016). TOP10 chemically competent or BL21 *E. coli* cells (Thermo Fisher, Cat# C404003) were transformed with the appropriate plasmid (see Supplementary Table 1 for plasmids used in this study) and grown to an OD600 of 0.6 in LB medium containing the appropriate antibiotic at 37 °C. Heterologous protein expression was then initiated with 0.02% w/v L-arabinose (Sigma A3256) or 0.5 mM IPTG overnight at 30 °C. Cells were collected by centrifugation (4000 g, 10 min) and stored at -80 °C. For purification, cells were resuspended and lysed in BugBuster Master Mix Protein Extraction Reagent (MilliporeSigma, Cat# 71456) at room temperature following the manufacturer’s recommendation. Lysate was clarified via centrifugation (15000 g, 10 min), and the soluble fraction of His-tagged proteins was purified using His SpinTrap columns (Cytiva, Cat# 28401353) following the manufacturer’s protocol. The eluate protein was buffer exchanged into PBS pH 7.4 and concentrated with 3 kDa Amicon Ultra-4 centrifugal filter units (MilliporeSigma, Cat# UFC800308). Proteins were frozen and stored at -80 °C. All purification steps were performed on ice unless otherwise noted.

All plasmids used for expression of fluorescent biosensors can be found in Supplementary Table 1.

### Fluorescence spectroscopy

Purified SED1 was added to a solution of 20 mM PBS pH 7.4 with and without 10% PEG4000 (w/v). Emission spectra of both solutions were collected at room temperature by exciting at 410 nm (5-nm slit width) and collecting emission from 425– 650 nm (2-nm slit width and a step size of 1 nm), with an exposure time of 0.1 s.

Purified MatryoshCaMP6s and jREX-GECO1 were added to a buffer solution with 35 µM Ca^2+^ and in the absence of Ca^2+^ (Invitrogen, Cat# C3008MP). MatryoshCaMP6s emission spectra were recorded by exciting at 440 nm (5-nm slit width) and collecting emission from 480–700 nm (1-nm slit width and a step size of 1 nm). The excitation spectra of jREX-GECO1 were recorded by exciting from 400–595 nm (1 nm slit width and a step size of 1 nm) and collecting emission at 630 nm (20 nm slit width) with an exposure time of 0.1 s.

Purified pHRed was added to buffers of pH 5.5 (50 mM MES-NaOH, 300 mM NaCl) and pH 9.0 (50 mM Tris-HCl, 300 mM NaCl) in a cuvette and loaded into a fluorimeter (Horiba Fluorolog 3). Excitation spectra of both solutions were collected by exciting from 400–590 nm (1-nm slit width and a step size of 1 nm) and monitoring emission at 630 nm (20-nm slit width), with an exposure time of 0.5 s. A 594 nm long-pass filter was added before the detector to reduce scattered light detection. Excitation spectra of the frozen samples were collected as above, except that the exposure time was changed to 0.1 s, and five spectra collected in succession were averaged to minimize the effect of any nitrogen boiling.

Purified pHmScarlet was added to buffers of pH 5.5 (50 mM MES-NaOH, 300 mM NaCl) and pH 8 (50 mM Tris-HCl, 300 mM NaCl). Excitation spectra of both solutions at RT and after freezing were collected with 400–585 nm excitation (1-nm slit width and a step size of 1 nm) and emission at 612 nm (20-nm slit width), with an exposure time of 0.1 s.

Purified pHluorin2 was added to buffers of pH 5.5 (50 mM MES-NaOH, 300 mM NaCl) and pH 7.5 (50 mM MOPS-HCl, 300 mM NaCl). Excitation spectra of both solutions at RT and freezing were collected with 350–500 nm excitation (1-nm slit width and a step size of 1 nm) and emission at 520 nm (20-nm slit width), with an exposure time of 0.1 s.

For cryogenic spectra collection, each room-temperature sensor sample was frozen in liquid nitrogen and held within a custom cryogenic sample holder in a transparent liquid nitrogen dewar. Cryogenic measurements were collected using the same conditions as at room temperature for each sensor unless noted.

### High-pressure freezing of purified biosensors

Purified pHRed and pHmScarlet were added to buffer solutions with varying pH. The buffers contained 300 mM NaCl and 50 mM of MES for pH 5.5–6.0, MOPS for pH 6.5– 8.0, and Tris for pH 8.5–9.0. Purified SED1 was added to a solution of 20 mM PBS pH 7.4 with various concentrations of PEG4000: 0, 2, 4, 8, 12, 16, and 20% (w/v). Purified jREX-GECO1 was added to a buffer solution at various Ca^2+^ concentrations. The solutions were made using a commercially available calcium calibration buffer kit (Invitrogen, Cat# C3008MP) at the following concentrations: 0, 0.015, 0.058, 0.135, 0.316, 0.542, 1.215, and 35.0 µM.

An aliquot of 3 µL or 10 µL of each sample was added to the 0.2 mm side of a glow-discharged Type A 3 mm aluminum or 6 mm gold-plated copper carrier, respectively, and sandwiched with the flat side of the corresponding Type B carrier. The sandwich carrier was immediately transferred to a high-pressure freezer (Leica EM ICE, Leica Microsystems) to generate a thick layer (∼0.2 mm) of frozen protein. After high-pressure freezing, the gold carriers were stored in liquid nitrogen.

### Production of bacterial strains and cell lines

#### Production of the jREX-GECO1 U2OS cell line

Human U2OS cells were maintained at 37 °C and 5% CO_2_ and cultured for no more than 20 passages in Dulbecco’s Modified Eagle Medium (DMEM) supplemented with 10% fetal bovine serum, Pen/Strep (Life Technologies, catalog number 15140122), and l-glutamine (Life Technologies, catalog number 25030081).

To generate jREX-GECO1 lentivirus for cell line production, 4 x 10^6^ HEK293T cells were seeded into 10 cm plates and were transfected the following day with 45 μL Mirus LT1 transfection reagent (MIR2300) added to a mixture of 5 μg each (15 μg total) of plasmids VSV-G (Addgene, Cat# 8454), R8.74 (Addgene, Cat# 22036), and pLV-JREX-GECO1-IRES-puro (this study) in 1.5 mL Opti-MEM (Thermo Fisher, Cat# 31985062) according to the manufacturer’s recommendations. Plasmid sequences were confirmed by full plasmid sequencing. Virus-containing supernatant was obtained 3 d post-transfection, clarified by centrifugation at 2,000 rpm for 2 min, and concentrated 10-fold using Lenti-X concentrator (Takara Bio, Cat# 631231). On the day prior to transduction, U2OS cells were seeded at a density of 1 x 10^5^ cells per well into individual wells of a 12-well plate (Cat# 07–200-82). Stable pools of cells expressing the desired protein of interest were obtained by titrating virus concentrate to achieve near 100% expression efficiency and selected with 2 μg/mL puromycin (Fisher, Cat# AAJ67236XF) for two weeks before conducting experiments.

#### Production of CbbS-pHmScarlet H. neapolitanus strain

Construct cloning and transformation of *H. neapolitanus* were performed as described previously (MacCready et al., 2021). Briefly, the fusion constructs were generated using Gibson Assembly, verified by sequencing, and transformed into competent cells of *H. neapolitanus* (*ATCC® 23641™*) for homologous recombination into the native carboxysome (*cso*) operon. The sequence encoding the fluorescent protein was attached to the 3’ region of the native *cbbS* coding sequence, separated by a GSGSGS linker. A kanamycin resistance cassette was inserted immediately downstream of the *cbbS* coding sequence. Transformants were selected by plating on S-6 medium with kanamycin (50 µg/mL). Colonies were restreaked, and transformants were verified for insertion by PCR and fluorescence microscopy. Colonies positive for the fusion and negative for the plasmid were grown in S-6 medium and stored at -80 °C in 10% DMSO.

All strains used in this study can be found in Supplementary Table 2.

### Cell growth and sample preparation

#### Sample preparation of S. cerevisiae

*S. cerevisiae* cells expressing SED1 were streaked onto a YNB SD-ura plate and incubated at 30 °C for 2–3 days. Single colonies were inoculated in 5 mL of YNB SD-ura medium and grown for 18–20 h at 30 °C with shaking at 250 RPM. An aliquot of 1 mL of cells was harvested by centrifugation and resuspended in 100 µL of spent YNB SD-ura medium. For osmotic shock, 1 M NaCl was added to 50 µL of the cell resuspension approximately 10 min before plunge freezing. A 3-µL aliquot of each sample was loaded on a glow-discharged (30 s at 25 mA) carbon finder grid (EMS, Cat# cf-200-cu-50), back-blotted for 2 s, and plunge-frozen in liquid ethane (Gatan CP3 or Leica GP).

#### Sample preparation of U2OS cells

Gold Quantifoil R2/2 200 mesh London Finder EM grids (Sigma, Cat# TEM-LFH2100AR1.3) were glow-discharged for 30 s at 25 mA. After glow discharging, 100 µL of DMEM was added to the bottom of the central 4 wells of an eight-chamber slide with removable wells (Sigma, Cat# PEZGS0816), and EM grids were added to the bottom of each of these wells in a laminar flow hood. JREX-GECO1-expressing U2OS cells were split using 0.05% trypsin in PBS (Fisher Scientific, Cat# 25-200-056) and resuspended to 1 x 10^5^ cells/mL. 200 µL of cells was transferred to each well atop a single EM grid, yielding a density of 2 x 10^4^ cells/grid. Cells on grids were left to recover overnight.

The following day, grids were removed from the multi-well plate and placed into the chamber of a pre-equilibrated Leica GP plunger at 37 °C and 90% humidity. Cells were then incubated for 3 min in the chamber with a solution containing the membrane pore-forming protein Streptolysin O (SLO) (Sigma, Cat# S5265-25KU) sufficient to permeabilize 90% of HeLa cells, as used previously (Teng et al., 2016). Cells were then plunged immediately into liquid ethane after back-blotting for 4 s and moved to liquid nitrogen for storage.

#### Sample preparation of H. neapolitanus

Frozen *H. neapolitanus* cells were streaked onto an S-6 plate and incubated for 4 days at 30 °C in air supplemented with 5% CO_2_. Single colonies were then inoculated in 6 mL of S-6 medium and incubated for 32 h at 30 °C with shaking at 130 RPM in air supplemented with 5% CO_2_. This culture was diluted (1:100) into 6 mL of fresh S-6 medium and grown for 20–24 h. Cells were harvested by centrifugation at 5,000 rpm for 5 min and resuspended in 20 µL of spent S-6 medium. An 18-µL aliquot of the resuspension was mixed with 2 µL of a solution of 5x concentrated 15-nm gold beads (EMS, Cat# 25489) resuspended in S-6 medium and loaded on glow-discharged (30 s at 15 mA) finder grids (Quantifoil on H2 London Finder Grid), back-blotted for 5 s, and plunge frozen in liquid ethane (Leica GP).

### Cryogenic fluorescence microscopy

Cryogenic fluorescence microscopy of purified SED1, pHRed, and jREX-GECO1 and of SED1-expressing *S. cerevisiae* cells was performed at 77 K on a home-built open cup cryostat described previously (Perez et al., 2024; Perez, 2024). Liquid nitrogen was added to an insulated stainless-steel cup, cooling a mounted copper pedestal to 77 K. The sample planchette or grid was then loaded onto the pedestal. The samples were imaged with a Nikon CFI Plan Achromat LWD IMSI 100 XC (NA 0.85) through a 1 mm glass window. The appropriate excitation and emission filters (described below) were used for each biosensor, and emission was collected by an EMCCD (Andor iXon Ultra 897). Frozen SED1 was excited with 425 nm light (Omicron LuxX) passing through a 427/10 nm excitation filter and with a peak intensity of 19.2 W/cm^2^ at the sample plane. Fluorescence emission over the range 450-565 nm was distinguished from scattered excitation by a dichroic mirror, a 440 nm long-pass filter, and two bandpass filters for CFP and YFP emission collection: a 465/30 nm bandpass filter for 3–4 s and a 545/40 nm bandpass filter for 3–4 s, respectively. Images were collected at 1 Hz with a gain setting of 3. Measurements were repeated across at least 5 regions on the carrier. Plunge-frozen *S. cerevisiae* cells were imaged with the same optical filters as purified SED1 but with a 425 nm excitation intensity of 12.6 W/cm^2^ and a gain setting of 10. Frozen pHRed was excited with interleaved 425 nm (Omicron LuxX; 8.3 W/cm^2^) and 561 nm (Crystalaser; 0.38 W/cm^2^). Fluorescence emission was filtered with a dichroic mirror (Chroma 59003bs) and a 594 nm long-pass filter (Semrock). Images were collected at 10 Hz with a gain setting of 300. Frozen jREX-GECO1 was excited with 488-nm light (Coherent Sapphire; 12.6 W/cm^2^) for 3–4 s and subsequently excited with 561-nm light (Crystalaser; 12.8 W/cm^2^) for 3–4 s. Fluorescence emission was filtered with a dichroic mirror (Chroma ZT405/488/561rpc) and a bandpass filter (Semrock FF01-607/70-25). Images were collected at 1 Hz with a gain setting of 3. A white-light image was collected after all measurements for image registration.

Cryogenic fluorescence microscopy of purified pHmScarlet and CbbS-pHmScarlet in *H. neapolitanus* cells was performed at 77 K using a similar home-built open-cup cryostat. Liquid nitrogen was added to an insulated stainless-steel cup, cooling a mounted aluminum alloy pedestal to 77 K. Samples were imaged with a room-temperature Nikon CFI S Plan Fluor ELWD 60XC (NA 0.7) through a 1-mm glass window. Emission was collected by a Hamamatsu Orca-Fusion BT Digital CMOS camera. Initial calibration of frozen pHmScarlet was performed using interleaved excitation with 488-nm light (Coherent; 5.73 W/cm^2^) and 561-nm light (Coherent; 0.3 W/cm^2^) with a 1-s period. Fluorescence emission was filtered with a 561-nm dichroic mirror (BrightLine Di03-561-t1) and a 561-nm longpass filter (EdgeBasic BLP02-561R-25). Images were collected at 10 Hz. *H. neapolitanus* cells expressing CbbS-pHmScarlet were imaged using interleaved excitation with 488-nm light (3.97 W/cm^2^) and 561-nm light (4.5 W/cm^2^) with a 2 s period. Images were collected at 5 Hz. Frozen pHmScarlet calibration samples at pH 5.5, 7.0, and 8.5 were imaged using these excitation intensities and interleaved with a period of 0.5 s. Images were collected at 100 Hz.

Ratiometric imaging of frozen jREX-GECO1 and vitrified U2OS cells was performed on the integrated Fluorescence Light Microscope (iFLM) unit of the Aquilos2 cryo-FIB (Thermo Scientific). Calibration samples were loaded onto the Aquilos2 stage using a 3 mm HPF-Carrier Shuttle (Thermo Scientific) and imaged with the 470-nm (50 ms at 5%) and 565 nm (50 ms at 5%) channels. U2OS samples were imaged with the 470-nm (2.5 s at 5%) and 565-nm (2.5 s at 5%) channels. A reflection image was collected post FIB-milling for image registration.

### Ratiometric calibration

Ratiometric calibration images were analyzed with a custom Python script. The average intensity of a 100 x 100-pixel region at the center of the image was recorded across the entire frame stack. The average intensity across the frames corresponding to each excitation wavelength or emission channel was averaged, except for pHRed. Because pHRed rapidly photobleached, only the first frame for each excitation was used for subsequent analysis. The average intensity for each wavelength or channel was then background-corrected by subtracting the average intensity of a blank high-pressure frozen sample containing buffer without fluorescent protein. The fluorescence ratio (F_ratio_) was then calculated by dividing one wavelength or channel background-corrected average intensity by the other. The average F_ratio_ across technical replicate measurements was then plotted for each environmental condition and fit to a continuous log function (Equation 1) by least-squares weighted by the associated error of each data point. Fit parameters for each biosensor can be found in Supplementary Table 3.

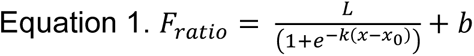

### Ratiometric analysis of cellular samples

F_ratio_ images of cell samples were generated with a series of custom Python scripts. First, the frames corresponding to each color channel were averaged. Afterward, a rough ratiometric image was calculated by taking the ratio between the two-channel average images. This rough ratiometric image was then used to perform cell (*S. cerevisiae* and *H. neapolitanus*) or grid square (U2OS) segmentation with the Python package Cellpose (Stringer et al., 2021). The segmentations were used to generate masks for subsequent analysis. The average channel images were then background-corrected using two procedures. First, a rolling ball background subtraction routine was applied, followed by a threshold-based background subtraction. A histogram of the pixel intensities reveals several peaks, with the lowest intensity peak corresponding to the background regions with only support film and no biological material. This peak was first isolated by taking the lowest 20% of the pixel intensities post rolling ball filter and then fit to a Gaussian distribution. The final offset was defined as the mean of the Gaussian fit plus two standard deviations. After background correction, the segmentation masks were used to define where the ratio between the two images was calculated. To generate the Ca^2+^ and pH maps, a pixel-wise model inversion was applied to corresponding F_ratio_ images. The inverse of the log function models derived from the calibration measurements was applied to each pixel in the F_ratio_ image.

### Focused-ion beam milling

*S. cerevisiae* and U2OS cell grids were loaded into an Aquilos2 (Thermo Fisher Scientific) for cryo-FIB milling. U2OS grids were screened using the iFLM as described above. SEM images were then collected to select lamella sites based on ratiometric fluorescence images. Sputter coating was performed for 15 s before and after GIS coating. Auto-TEM was then used to produce a lamella of approximately 200 nm thickness at each site. Rough milling was performed at 1.0 nA to achieve a thickness of approximately 1 µm, followed by a series of medium and fine milling steps at 0.5, 0.3, and 0.1 nA. The final polishing step was completed with 30–50 pA to produce lamella with a thickness of 170–220 nm. Grids were then retrieved from the Aquilos2 and stored in liquid nitrogen.

### Cryogenic electron tomography

Grids were loaded into a 300 kV Titan Krios (Thermo Fisher Scientific) equipped with a BioQuantum K3 Imaging Filter and a K3 Camera (Gatan) or a Selectris Imaging Filter and a Falcon 4i direct detector. Tomography of *S. cerevisiae* lamellae was performed by tilting the lamella between ± 48 ° in the electron beam using a dose-symmetric tilt scheme with a 2° tilt step. The unbinned pixel size was 4.625 Å, and the total dose was ∼120 e/Å^2^ collected at 19000x magnification with an 8-µm target defocus. Tomography of U2OS lamellae was performed by tilting the lamella between ± 60 ° in the electron beam using a dose-symmetric tilt scheme with a 3° tilt step. The unbinned pixel size was 2.68 Å, and the total dose was ∼120 e/Å^2^ collected at 33000x magnification with a 4 µm target defocus. Tomography of *H. neapolitanus* was performed by tilting the grid between ± 50° in the electron beam using a dose-symmetric tilt scheme with a 2.5° tilt step. The unbinned pixel size was 4.75 Å, and the total dose was ∼100 e/Å^2^ collected at 26000x magnification with a 20 µm target defocus.

### Tomogram image processing

*S. cerevisiae* and *H. neapolitanus* tomograms were reconstructed using the Etomo package in IMOD (Mastronarde and Held, 2017). Tilt series alignment was achieved using a selection of gold and non-gold (small ice particles) fiducials located in the periphery of *H. neapolitanus* cells or on *S. cerevisiae* lamellae, respectively. U2OS tomograms were processed within a Scipion environment (Conesa et al., 2023). Motion correction was performed using MotionCor2 (Zheng et al., 2017). CTFfind 5 was used for CTF estimation, and the tilt series were aligned and reconstructed using AreTomo2 (Zheng et al., 2022), followed by denoising in cryo-CARE and 3DEM (Buchholz et al., 2019). All tomograms were binned by 4 and annotated using MemBrain (Lamm et al., 2022) with default parameters to detect membranes and Dragonfly 3D (v2024.1) (Heebner et al., 2022) with a custom-trained neural network to segment ribosomes, cell wall, carboxysome shells, and the carboxysome lumen. All segmentations were then manually cleaned within Dragonfly 3D before exporting to ChimeraX for visualization.

### Electron and fluorescence microscopy image registration

Electron and fluorescence images were registered using a series of affine transformations. First, EM search map magnification images were manually aligned and stitched using the napari-stitcher package in Napari (Chiu et al., 2022). All subsequent processing was performed with a custom MATLAB script previously described (Dahlberg et al., 2020). For *S. cerevisiae* images, optical images were registered to an SEM image collected post-milling, using 15-20 grid holes to compute a projective transformation carrying the optical data into the EM space. This SEM image was then registered to the search map image by identifying unique features, such as holes and lamella deformities. The search map image was then registered with the cryoET space by identifying ice contamination and cellular features visible in both the search map and the z-projection of the cryoET tomogram reconstruction. From these control-point pairs, a similarity transformation was computed to carry the search map image to the cryoET space. Sequential application of the projective and similarity transformations to the optical data carries the ratiometric fluorescence images into the cryoET space. For U2OS, the same computational workflow was applied to the following images in this order: pre-milling fluorescence, post-milling reflection, stitched EM atlas, overview, stitched search map, and z-projection of reconstructed tomogram. *H. neapolitanus* registration was achieved with the following images: fluorescence micrograph, EM overview, EM search map, and z-projection of reconstructed tomogram, using gold beads as fiducials. Fluorescence ratio, pH maps, and calcium maps were registered by applying the transformations sequentially. All registered overlays were produced using a custom Python script.

## Data Availability

All data supporting the findings of this study are included in the manuscript and its supplementary materials. Raw fluorescence micrographs and electron tomography reconstructions will be made available at the Stanford Digital Repository (doi.org/10.25740/wd385cn5299) and Electron Microscopy Data Bank at a later date. Additional data are available from the corresponding author upon request.

## Code Availability

All code used in this study is publicly available and can be found at: github.com/azaldegui/biosensor_CLEM

## Acknowledgements

We thank all members of the Dahlberg Lab for helpful discussions and advice throughout the project. We thank Drs. Bharti Singal and David Fernandez Martinez at the Stanford Cryo Electron Microscopy Centre (cEMc) for instrument maintenance and assistance. We thank Prof. W.E. Moerner for access to his lab space and cryogenic light microscope during the early stages of this work. We thank Prof. Soichi Wakatsuki for access to his lab space for protein purification. pRsetB-pHRed was a gift from Gary Yellen. RpHluorin2 was a gift from Geert van den Bogaart. pBAD-pHScarlet and pTorPE-jREX-GECO1 were a gift from Robert Campbell. pET15b-eGO-Matryoshka-LE-LP was a gift from Wolf Frommer. pCMV-VSV-G was a gift from Bob Weinberg. J.S.B. acknowledges support from the National Institutes of Health grant R01-GM144731. A.G.V. acknowledges support from the National Institutes of Health grant R35-GM152128. P.D.D. and C.A.A. were supported by the U.S. Department of Energy, Office of Science, Office of Biological and Environmental Research under Award Number DE-AC02-76SF00515 (FWP 100883).

## Author contributions

C.A.A. and P.D.D. conceptualized the project. C.A.A., J.A.S., and C.A.M. purified proteins. C.A.A., D.P., and P.D.D. performed spectroscopy and light microscopy. C.A.A. and K.M.R. generated and cultured strains. C.A.A., K.M.R., and P.D.D. performed cryoFIB and cryoET. C.A.A., K.M.R., and W.P.D. analyzed cryoET data. C.A.A. analyzed light microscopy data. P.D.D., J.R.D., J.S.B., and A.G.V. provided funding and project supervision. C.A.A. wrote the manuscript with input from other authors.

