## Supplementary Figures for "Fluorescent Biosensors for Cryogenic Correlative Light and Electron Microscopy"

**Supplementary Table 1. Plasmids used in this study**

| Plasmid | Description | Source |
|---|---|---|
| *pDEST-HisMBP-SED1* | Bacterial expression of SED1 | Addgene #178191, [1] |
| *pRsetB-pHRed* | Bacterial expression of pHRed | Addgene #31472, [2] |
| *RpHluorin2* | Bacterial expression of pHluorin2 | Addgene #180227, [3] |
| *pBAD-pHScarlet* | Bacterial expression of pHmScarlet | Addgene #191258 |
| *pET15b-eGO-Matryoshka-LE-LP* | Bacterial expression of MatryoshCaMP6s | Addgene #100017, [4] |
| *pTorPE-jREX-GECO1* | Bacterial expression of jREX-GECO1 | Addgene #113941, [5] |
| *pLV-JREX-GECO1-IRES-puro* | For stable expression of jREX-GECO1 into U2OS cells | This study |
| *pCMV-VSV-G* | For transfection of jREX-GECO1 gene into U2OS cells | Addgene #8454, [6] |
| *cbbS-pHmScarlet* | For chromosomal replacement of the native *cbbS* gene in *H. neapolitanus* | This study |

**Supplementary Table 2. Strains used in this study**

| **Strain** | **Description** | **Source** |
|---|---|---|
| *S. cerevisiae* (SED1) | *S. cerevisiae* (BY4742) with *pDRFLIP38-SED1* | [1] |
| AGV_Hn_1 | *H. neapolitanus* | ATCC |
| AGV_Hn_547 | *H. neapolitanus cbbS::cbbS-pHmScarlet* | This study |

**Supplementary Table 3. Fit parameters for biosensor calibrations**

| **Biosensor** | **Figure** | **L** | **k** | $\mathbf{x_0}$ | **b** |
|---|---|---|---|---|---|
| SED1 | Main. 2c | 4.35 | 0.59 | 8.47 | 2.73 |
| pHRed | SI. 5b | 9.04 | 1.27 | 8.17 | 1.6 |
| pHmScarlet | Main. 2i | 102.13 | -2.93 | 6.54 | 1.48 |
| pHmScarlet | SI. 8a | 5.72 | -2.93 | 6.54 | 0.04 |
| jREX-GECO1 | Main. 2f | 2.4 | 2 | 0.15 | -1.07 |
| jREX-GECO1 | SI. 7a | 1.68 | 1.34 | 0.97 | 0 |

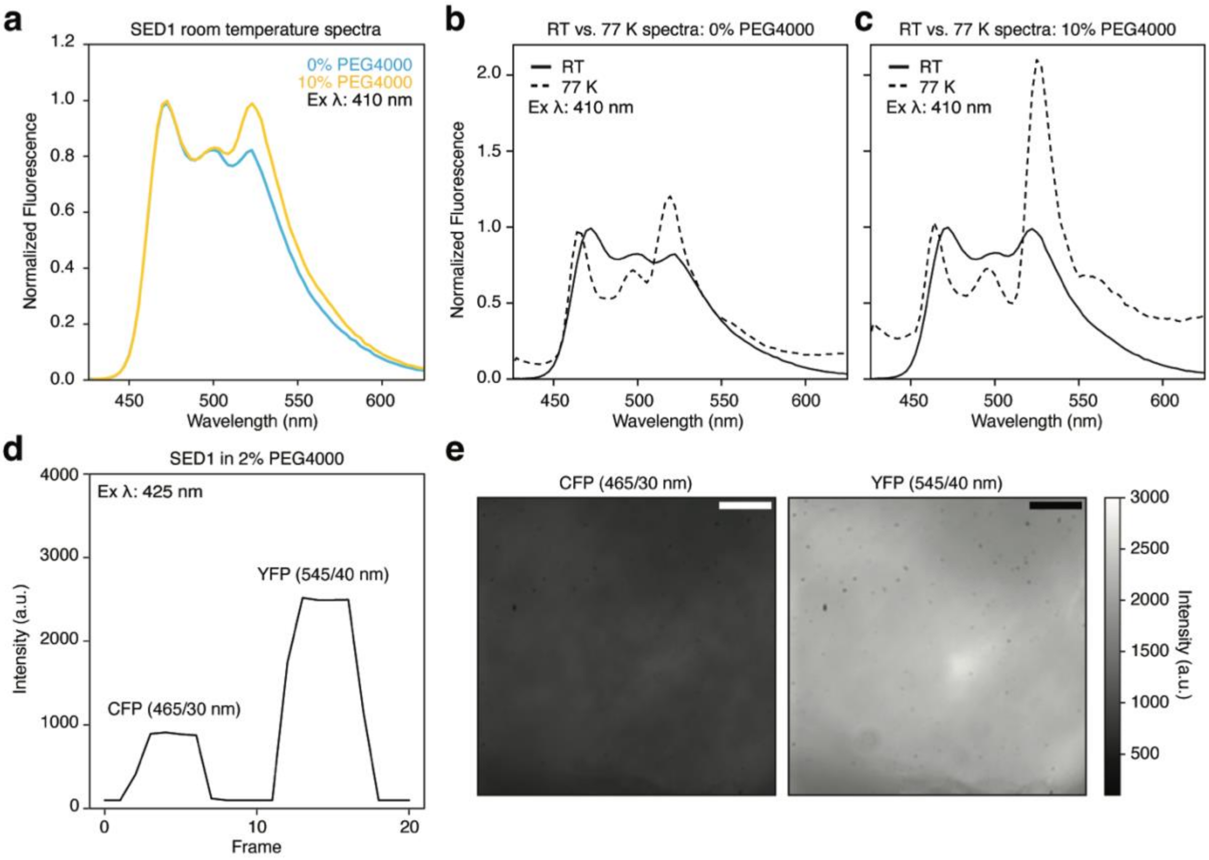


**Supplementary Figure 1: *In vitro* characterization of SED1.**
**a)** Fluorescence emission spectra of SED1 in the presence (yellow) and absence (blue) of 10% PEG4000 (w/v) at room temperature (RT). **b)** Fluorescence emission spectra of SED1 in 0% PEG4000 at RT (solid) and 77 K (dashed). **c)** Fluorescence emission spectra of SED1 in 10% PEG4000 at RT (solid) and 77 K (dashed). Emission spectra are normalized to their respective mCerulean peak at 465 nm. **d)** Representative fluorescence intensity trace of high-pressure frozen SED1 in 2% PEG4000 using CFP (465/30 nm) and YFP (545/40 nm) emission filters. **e)** Representative average fluorescent images of high-pressure frozen SED1 in 2% PEG4000. Scale bars: 15 µm.

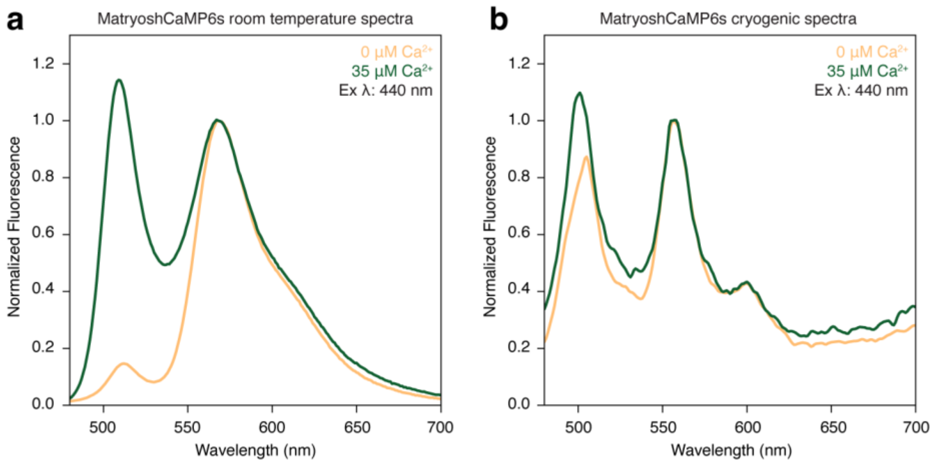


**Supplementary Figure 2: Emission spectra of MatryoshCaMP6s at room and cryogenic temperatures.**

**a)** Fluorescence emission spectra of MatryoshCaMP6s in the presence and absence of 35 µM $Ca^{2+}$ at room temperature. Emission spectra are normalized to their respective LSS-mOrange peak at 570 nm. **b)** As in **a**, but at 77 K.

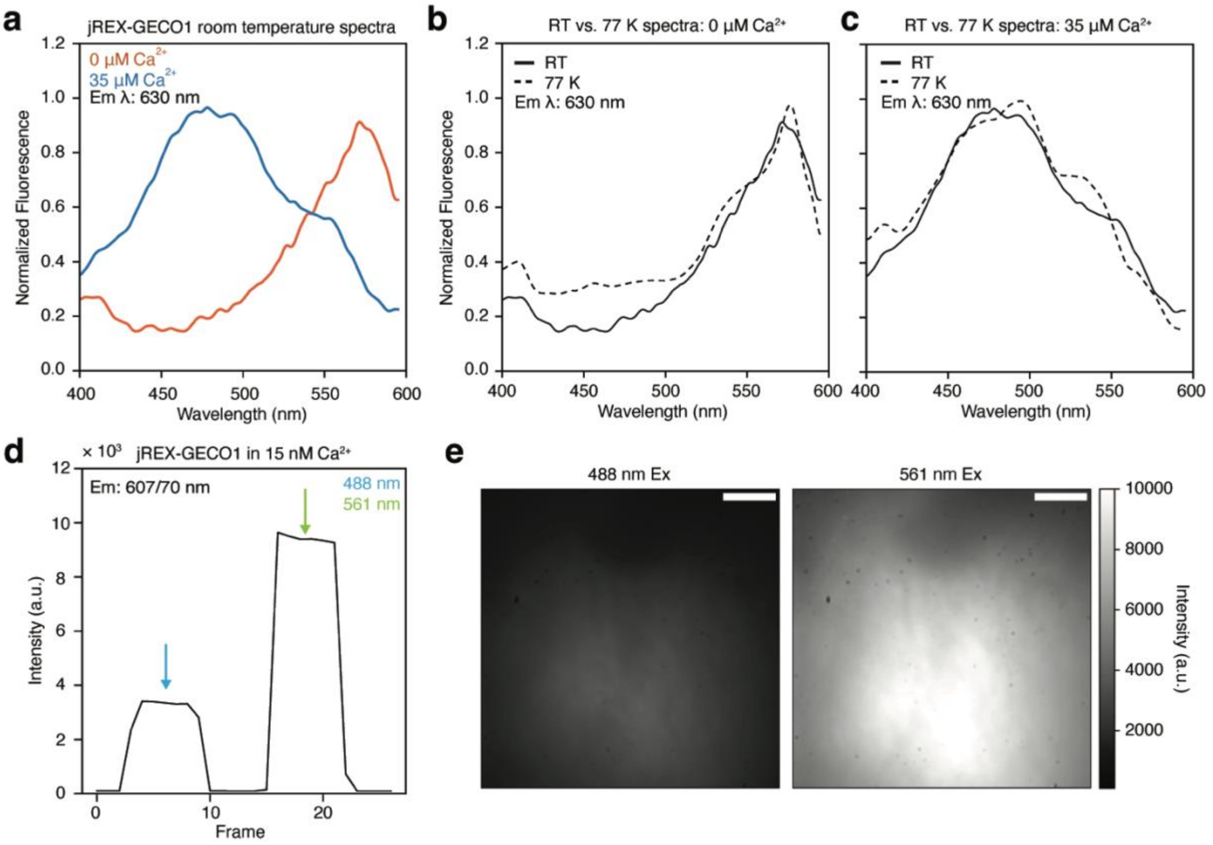


**Supplementary Figure 3: *In vitro* characterization of jREX-GECO1. a)** Room-temperature excitation spectra of jREX-GECO1 in the presence and absence of $Ca^{2+}$ (35 μM). **b)** Fluorescence excitation spectra of jREX-GECO1 in the absence of $Ca^{2+}$ at RT (solid) and 77 K (dashed). **c)** Fluorescence excitation spectra of jREX-GECO1 in the presence of 35 μM $Ca^{2+}$ at RT (solid) and 77 K (dashed). Excitation spectra are normalized independently. **d)** Representative fluorescence intensity trace of high-pressure frozen jREX-GECO1 in 0.015 nM $Ca^{2+}$ using 488 nm and 561 nm excitation lasers. **e)** Representative average fluorescent images of high-pressure frozen jREX-GECO1 in 15 nM $Ca^{2}$. Scale bars: 15 μm.

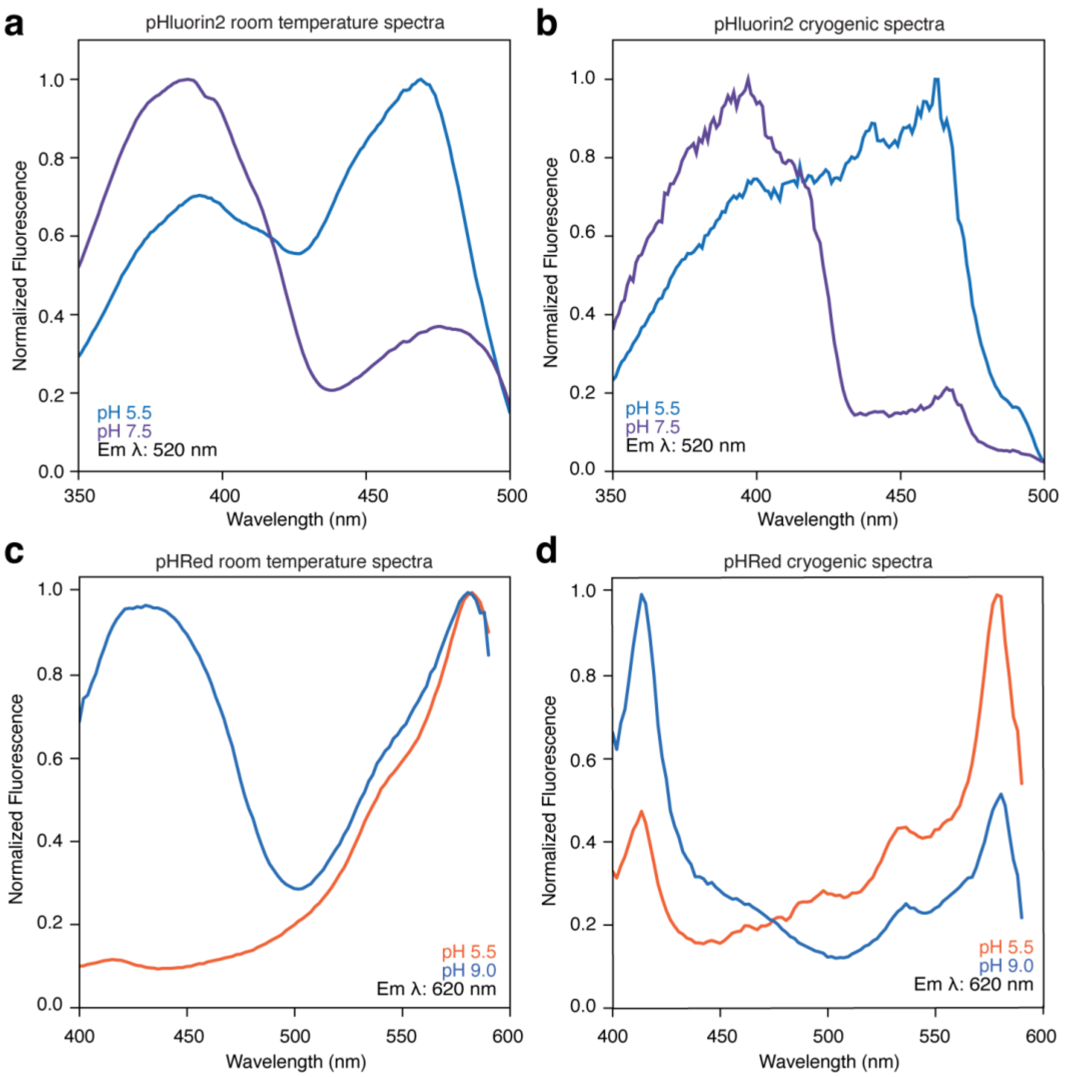


**Supplementary Figure 4: Fluorescence spectra of pH sensors at room and cryogenic temperatures. a)** Room-temperature excitation spectra of pHluorin2 at pH 5.5 and pH 7.5. **b)** As in **a**, but at 77 K. **c)** Room-temperature excitation spectra of pHRed at pH 5.5 and pH 9.0. **d)** As in **c**, but at 77 K. Excitation spectra are normalized independently.

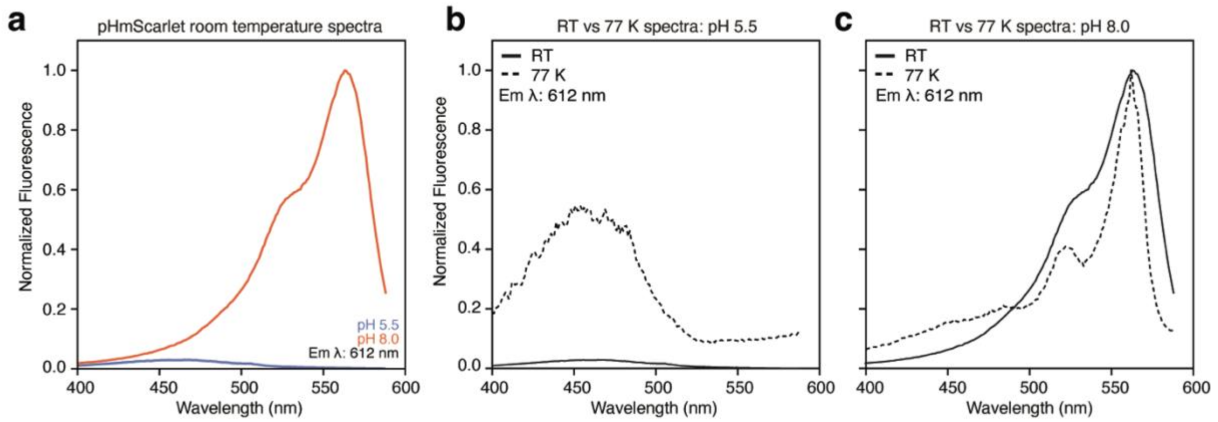


**Supplementary Figure 5: *In vitro* characterization of pHmScarlet. a)** Room-temperature excitation spectra of pHmScarlet at pH 5.5 (blue) and pH 8.0 (orange). **b)** Fluorescence excitation spectra of pHmScarlet at pH 5.5 at RT (solid) and 77 K (dashed). **c)** Fluorescence excitation spectra of pHmScarlet at pH 8.0 at RT (solid) and 77 K (dashed). Excitation spectra are normalized to the maximum of the corresponding pH 8 spectrum.

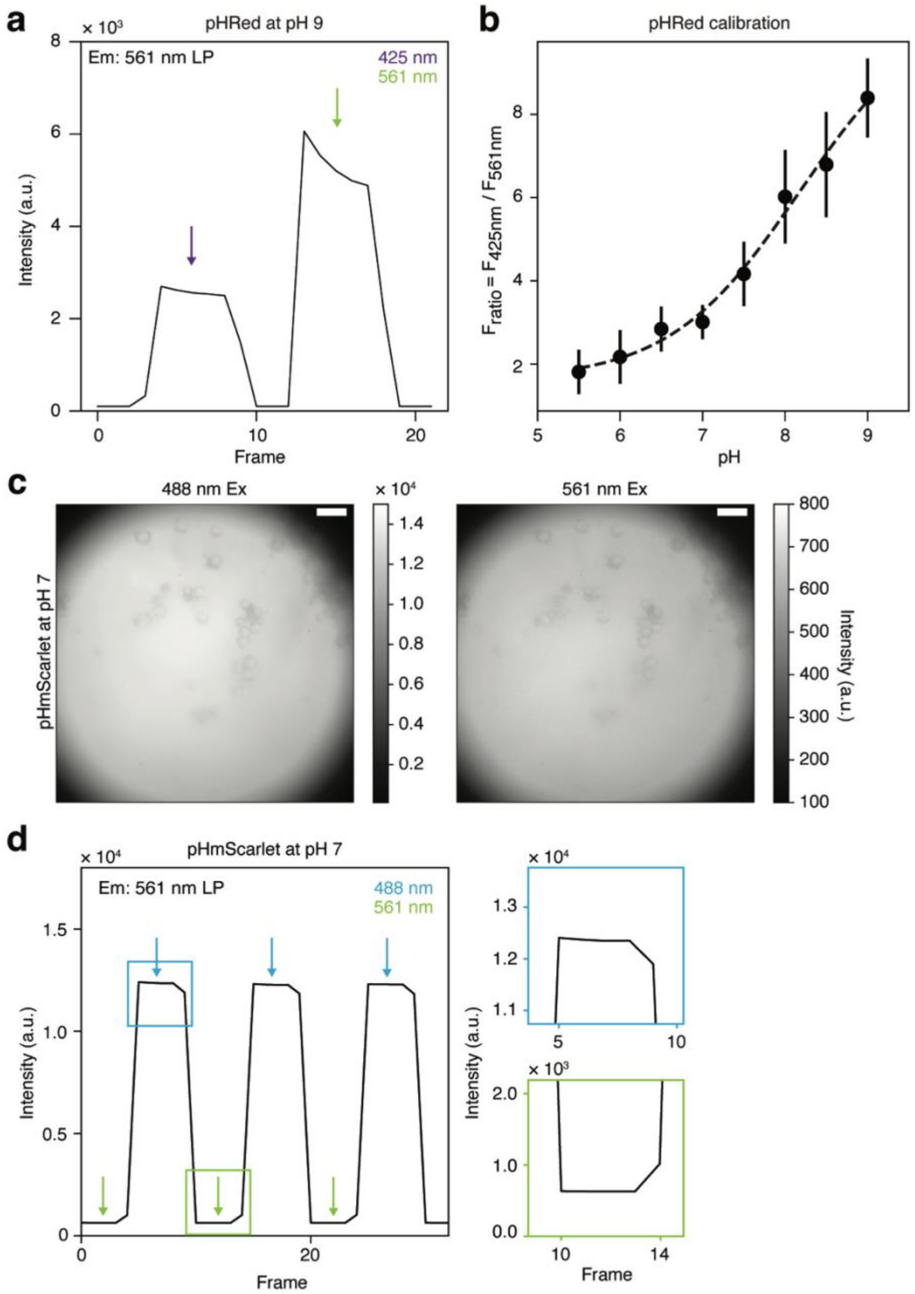


**Supplementary Figure 6: pHRed intensity trace and calibration.**
**a)** Fluorescence intensity trace of high-pressure frozen pHRed at pH 9. Intensity peaks are labeled with their respective laser excitation wavelengths. 561 nm excitation results in rapid photobleaching of pHRed. **b)** $F_{ratio}$ for 425- and 561-nm excitation of high-pressure frozen pHRed as a function of pH. Points and error bars represent the average and standard deviation of at least 5 ratiometric measurements for each condition. The dashed line is the fit to a continuous logistic function. **c)** Representative average fluorescent images of high-pressure frozen pHmScarlet in pH 7.0. Scale bars: 25 µm. **d)** Representative fluorescence intensity trace of high-pressure frozen pHmScarlet at pH 7 using 488 nm and 561 nm excitation lasers. Blue inset is zoom into 488 nm excitation trace. Green inset is zoom into 561 nm excitation trace.

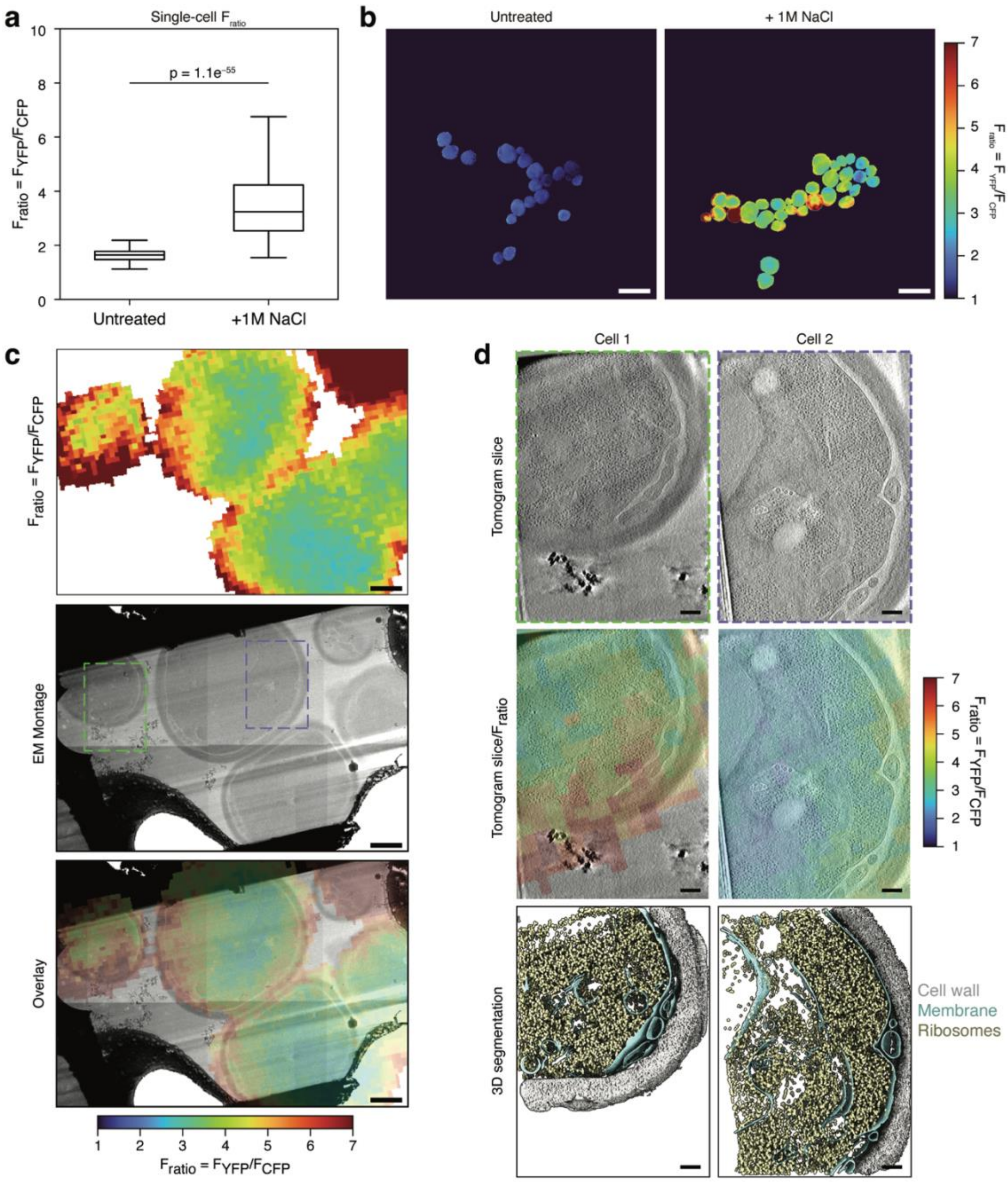


**Supplementary Figure 7: CryoCLEM of SED1 in *S. cerevisiae***

**a)** Single-cell average $F_{ratio}$ with ($N$ = 267 cells) and without ($N$ = 55 cells) osmotic shock. **b)** $F_{ratio}$ (YFP/CFP) fluorescence images of cells with and without osmotic shock. Scale bars: 10 µm. **c)** The top panel is a registered $F_{ratio}$ image corresponding to an EM montage of a lamella of osmotically shocked cells (middle panel). The bottom panel shows the overlay of the montage and the $F_{ratio}$ image. Colored dashed boxes are the tomogram collection regions shown in d. Scale bars: 1 µm. **d)** The top row shows slices from the reconstructed tomograms of the regions shown in **c**. The middle row shows the tomogram slices overlaid with the $F_{ratio}$ image. The bottom row shows the segmentation of ribosomes (gold), membrane (teal), and cell wall (gray). Scale bars: 200 nm.

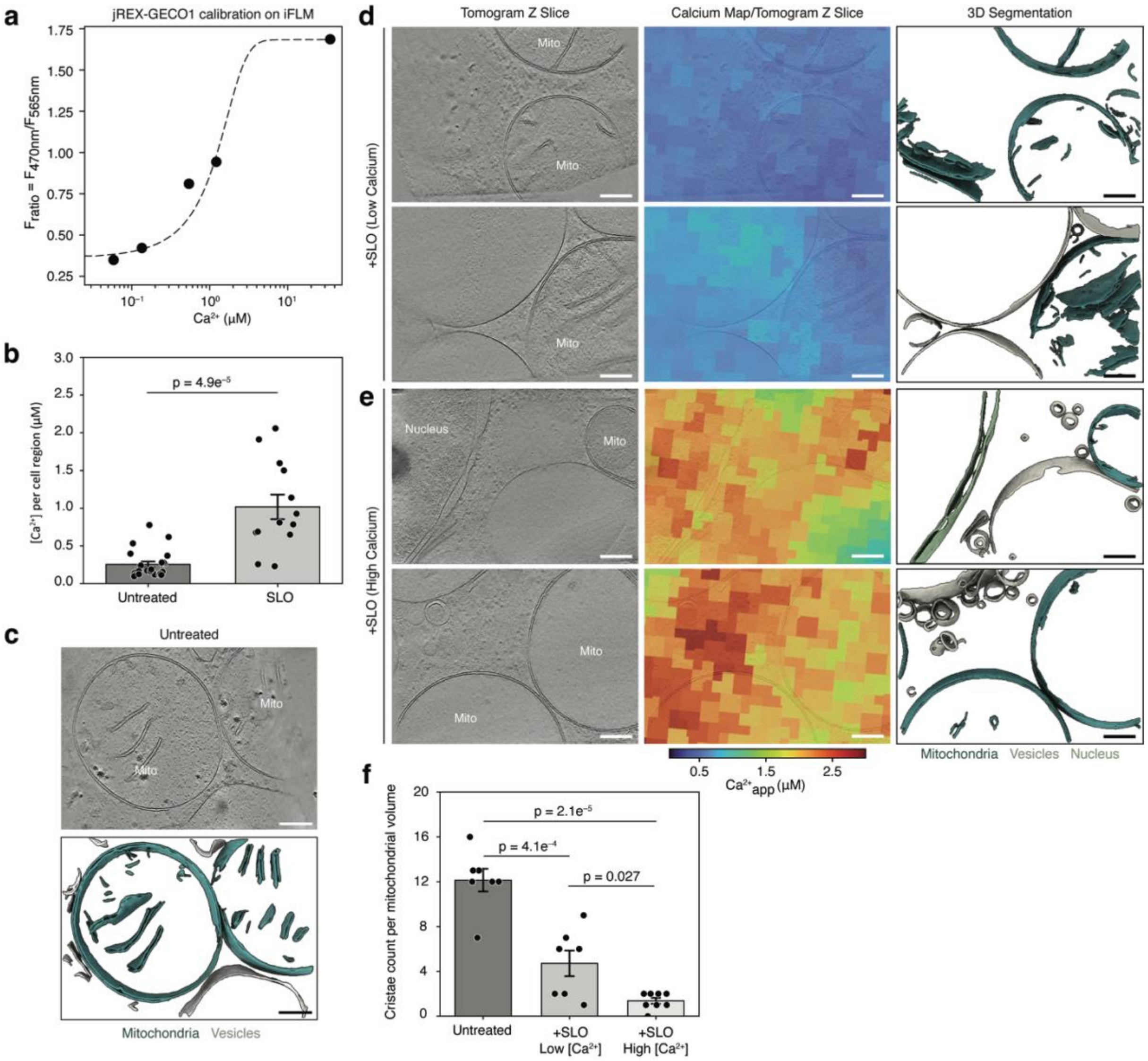


**Supplementary Figure 8: CryoCLEM of jREX-GECO1 in U2OS cells.**
**a)** $F_{ratio}$ for 470- and 565-nm excitation of high-pressure frozen jREX-GECO1 as a function of $Ca^{2+}$ concentration. The dataset was collected on the iFLM within an Aquilos2 system. The dashed line is the fit to a continuous logistic function. **b)** Average $Ca^{2+}$ concentration for untreated (*N* = 22 regions) and SLO-treated (N = 13 regions) U2OS cells. **c)** The top panel shows a z-slice from the reconstructed tomogram of an untreated U2OS cell. The bottom panel shows the segmentation of mitochondria (teal) and vesicles (gray). **d)** Two reconstructed tomogram z-slices are shown for SLO-treated U2OS cells with a low measured calcium level. The middle panels show the corresponding tomogram z-slices overlaid with the correlated calcium map. The right panels show the segmentation of mitochondria (teal), vesicles (gray), and the nuclear membrane (green). **e)** As in **d**, but for a cell with a high measured calcium level. Scale bars: 200 nm. **f)** Mitochondrial cristae counts categorized by sample and $Ca^{2+}$ concentration. *N* = 7 mitochondria from 4 to 7 tomograms for each group.

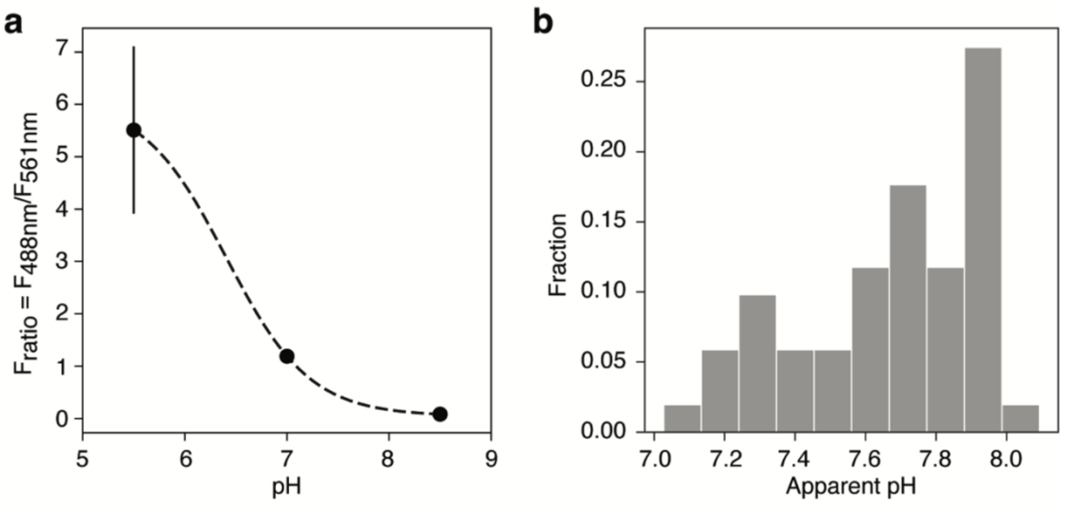


**Supplementary Figure 9: CryoCLEM of pHmScarlet in *H. neapolitanus*.**
**a)** $F_{ratio}$ for 488- and 561-nm excitation of high-pressure frozen pHmScarlet as a function of pH using the same laser excitation power densities used for cryogenic cellular imaging. Each calibration point represents the average of at least 5 ratiometric measurements for each condition. Error bars represent the standard deviation. The dashed line is the fit to a continuous logistic function derived from the calibration in Fig. 2i. This was a control to ensure that the significantly different excitation intensities used for imaging did not change the calibration. **b)** Distribution of the apparent pH from single *H. neapolitanus* cells measured using cryogenic ratiometric imaging of pHmScarlet. $N$ = 51 cells.
